# Genetic diversity in global populations of the Critically Endangered addax (*Addax nasomaculatus*) and its implications for conservation

**DOI:** 10.1101/2022.07.07.499131

**Authors:** Kara L Dicks, Alex D Ball, Lisa Banfield, Violeta Barrios, Mohamed Boufaroua, Abdelkader Chetoui, Justin Chuven, Mark Craig, Mohammed Yousef Al Faqeer, Hamissou Halilou Mallam Garba, Hela Guedara, Abdoulaye Harouna, Jamie Ivy, Chawki Najjar, Marie Petretto, Ricardo Pusey, Thomas Rabeil, Philip Riordan, Helen V Senn, Ezzedine Taghouti, Tim Wacher, Tim Woodfine, Tania Gilbert

**Author notes:** **Corresponding Author:** Kara L Dicks, RZSS WildGenes, Royal Zoological Society of Scotland, 134 Corstorphine Road, Edinburgh, EH12 6TS, United Kingdom.

## Abstract

Threatened species are frequently patchily distributed across small wild populations, *ex situ* populations managed with varying levels of intensity, and reintroduced populations. Best practice advocates for integrated management across *in situ* and *ex situ* populations. Wild addax (*Addax nasomaculatus*) now number fewer than 100 individuals, yet thousands of addax remain in *ex situ* populations, which can provide addax for reintroductions, as has been the case in Tunisia in the mid-1980s. However, integrated management requires genetic data to ascertain the relationships between wild and *ex situ* populations that have incomplete knowledge of founder origins, management histories and pedigrees. We undertook a global assessment of genetic diversity across wild, *ex situ*, and reintroduced populations in Tunisia to assist conservation planning for this Critically Endangered species. We show that the remnant wild populations retain more mitochondrial haplotypes which are more evolutionarily diverse than the entirety of the *ex situ* populations across Europe, North America and the United Arab Emirates, and the reintroduced Tunisian population. Additionally, 1704 SNPs revealed that whilst population structure within the *ex situ* population is minimal, each population carries unique diversity. Finally, we show that careful selection of founders and subsequent genetic management is vital to ensure genetic diversity is provided to, and minimise drift and inbreeding within, reintroductions. Our results highlight a vital need to conserve the last remaining wild addax population, and we provide a genetic foundation for determining integrated conservation strategies to prevent extinction and optimise future reintroductions.

## Introduction

Genetic diversity is one of three fundamental components of biodiversity in the Convention on Biological Diversity (Laikre et al., 2010; Secretary of the Convention on Biological Diversity, 1992) and the post-2020 global biodiversity framework (Convention on Biological Diversity, 2020; Hoban et al., 2020). It provides the foundation for evolution, enabling species to adapt to changing environmental and disease conditions (Allendorf, 1986; Frankham et al., 2010) and is therefore essential to the long-term persistence of species and populations (Lachapelle et al., 2017; Russell Lande & Shannon, 1996; Willi & Hoffmann, 2009). Small, fragmented populations are intrinsically vulnerable to reduced genetic diversity through inbreeding and genetic drift and thus at risk of accumulating deleterious mutations and suffering reduced fitness (Crnokrak & Roff, 1999; Frankham, 1995; R. Lande, 1994; Russell Lande & Shannon, 1996; Spielman et al., 2004).

*Ex situ* populations can play an essential role in preventing extinction (McGowan et al., 2017) and best practise emphasises the benefits of developing integrated management plans for threatened species that cover all *ex situ* and *in situ* populations, including those reintroduced to their indigenous range (Traylor-Holzer et al., 2019). Management varies substantially amongst *ex situ* populations, from internationally co-ordinated pedigree-based population management programmes that aim to maximise retention of genetic diversity (Ballou & Lacy, 1995), to low intensity management of large, independent populations (Wildt et al., 2019), with levels of genetic diversity varying substantially between such populations (Gooley et al., 2020, 2022; Humble et al., 2020; Ogden et al., 2020).

Management plans are increasingly incorporating populations with varied management histories, often requiring genetic data to determine the values of different populations relative to each other (Hogg et al., 2017; Marshall et al., 1999; Ogden et al., 2020) and providing information on how they should be managed to achieve a joint goal of species recovery (Hoffmann et al., 2015). This is especially important for species that have substantially larger populations under low-intensity management than they have under high-intensity management, or than are present in the wild. In practice, a balance may need to be struck between availability of individuals for reintroduction, demographic objectives, and genetic quality.

The biodiversity of the Sahelo-Saharan region has suffered a catastrophic decline in megafauna (Brito et al., 2014; Durant et al., 2014), including the Critically Endangered antelope, the addax (*Addax nasomaculatus*), which is now close to extinction in the wild (Durant et al., 2014; IUCN SSC Antelope Specialist Group, 2020; Stabach et al., 2017). Once abundant and widespread in the Sahelo-Saharan region (Figure 1A), it suffered rapid and catastrophic declines due to hunting, habitat degradation, regional insecurity and, most recently, the impacts of oil exploration in its remaining habitat (IUCN SSC Antelope Specialist Group, 2016). The wild population now numbers less than 100, restricted to just 0.68% of its historical range (IUCN SSC Antelope Specialist Group, 2020) in one population in the Tin Toumma desert of Niger up to the border with Chad (IUCN SSC Antelope Specialist Group, 2016, 2020; Rabeil et al., 2016).

**Figure 1.**
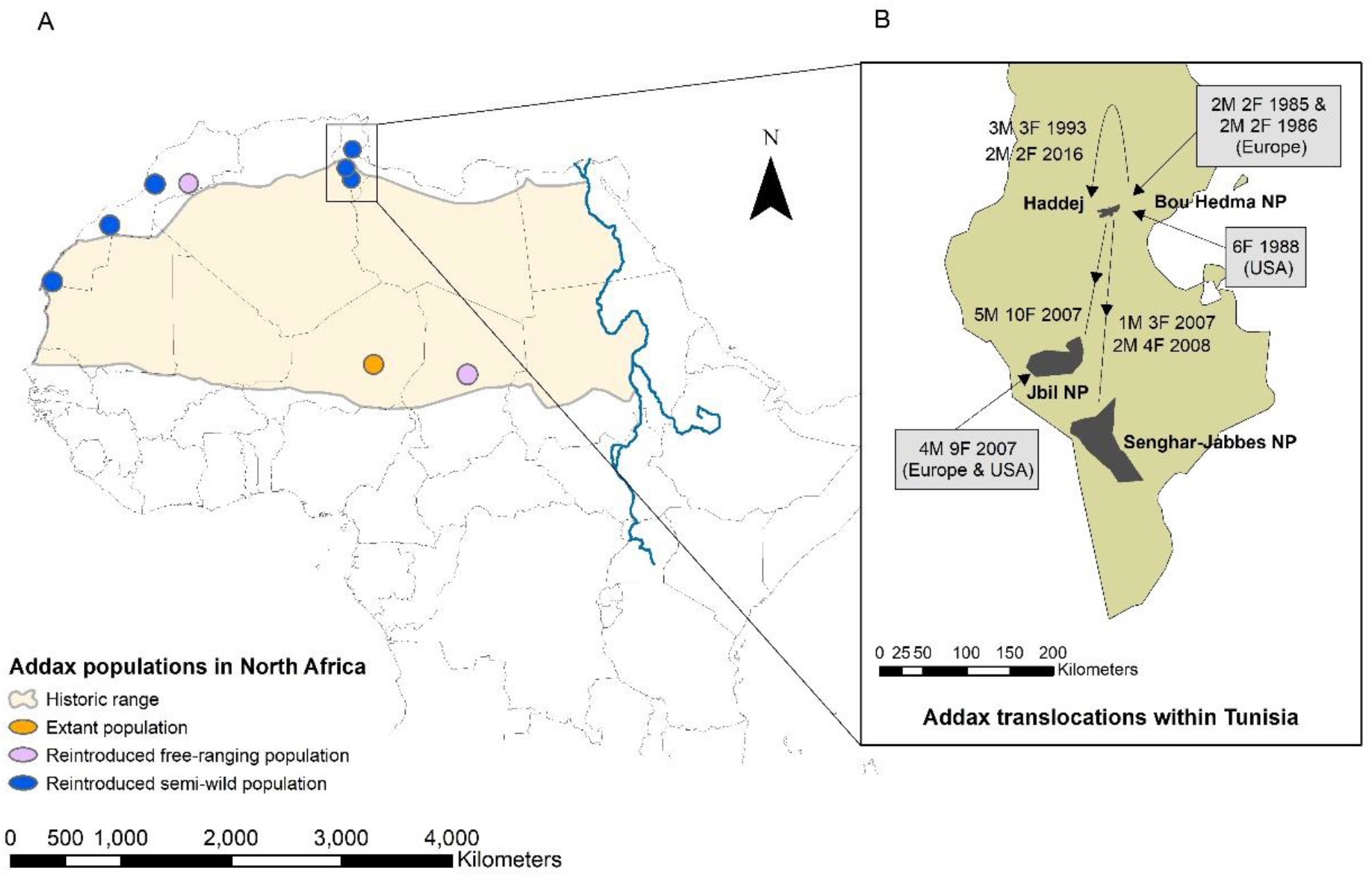
The historical range of addax (A) showing the historic addax range with locations of the remaining population in the Tin Toumma region, Niger, and free-ranging and semi-wild reintroduced populations. Inset (B) illustrates the reintroduction history of addax in Tunisia, indicating numbers of males (M) and females (F) translocated from each source population.

Whilst addax have been driven to near-extinction in the wild, the species has been maintained in global zoological institutions since 1920 (Krause, 2016). At the time of writing, there are nearly 1200 *ex situ* addax registered on the global ZIMS database (Species360, 2022) and several thousand unregistered addax in private populations in the USA and the Arabian Peninsula (IUCN SSC Antelope Specialist Group, 2016; Mallon & Chardonnet, 2020). The three largest regional populations registered in ZIMS are located in i) the United Arab Emirates (*N*=556), predominantly in the Al Ain Zoo (AAZ) (*N*=286) and Environment Agency – Abu Dhabi (EAD) facilities (*N*=142), ii) North America (*N*=256), and iii) Europe (*N*=212). The European and North American populations are managed within their respective co-ordinated population management programmes, the European Association of Zoos and Aquaria’s EAZA *Ex situ* Programme (EEP) and the Species Survival Plan® (SSP) under the Association of Zoos and Aquariums (AZA). As is typical for many species, founder histories are poorly known but founder numbers are thought to be relatively low for addax; the European studbook has 15 wild-caught founders listed (Krause, 2016) and the international studbook (incorporating the EEP and SSP) indicates up to a maximum of 42 founders (Enright, 2019), but it is likely to be far fewer.

Pedigrees and movements are poorly documented for most addax populations outside of these intensively managed programmes. Whilst anecdotal evidence hints towards the existence of novel founders, unmanaged populations in North America and the UAE are probably descended from many of the same wild-caught individuals as the EEP and SSP populations. The escalating biodiversity crisis in the Sahelo-Saharan region emphasises the importance of the *ex situ* addax population for conservation (Payne & Bro-Jørgensen, 2016), but a clear understanding of the genetic diversity and structure within the global addax population is crucial for formulating effective conservation action.

Relatively little is known about genetic diversity in either wild or *ex situ* populations of addax. A recent study by Hempel et al (2021) generated whole-genome sequencing data for an *ex situ* female and mitogenomes from 10 museum samples (1821 – 1926) originating from across the species’ historical range. These data indicated that, prior to recent declines in population numbers, there were relatively low levels of genetic diversity and minimal genetic structure, indicative of a large, highly panmictic population. Previous molecular genetic studies of *ex situ* addax populations are restricted to either single institutions (Armstrong et al., 2011) or regions (Ivy et al., 2016; Spevak et al., 1993), thus representing only a small proportion of the global population. Importantly, different methodologies are used across all studies, prohibiting comparison amongst populations.

Despite the lack of genetic information, *ex situ* populations have provided a source of animals for a series of historical conservation translocations to Tunisia (Bertram, 1988; Chardonnet, 2007; Correll & Houston, 1999; Gilbert et al., 2018), Morocco (Müller & Engel, 2004; SCF, 2020), and Chad (SCF, 2020). Re-establishing addax in former range countries began in Tunisia in 1985 with an introduction to Bou Hedma National Park (NP; with the entire population later transferred to the Haddej area within Bou Hedma NP) outside the indigenous range of the addax, but formed the first step to returning the species to a country that it had been absent from since 1932 (Gilbert et al., 2018). This population formed the basis for two reintroductions to Jbil and Senghar-Jabbes NPs, within their indigenous range, in the mid-2000s (Figure 1B). A single augmentation was carried out to Jbil NP in 2007 with addax carefully selected from the EEP and SSP to maximise genetic diversity. Although populations sizes of these population are monitored there has not been an evaluation of their genetics.

With addax on the brink of extinction in the wild and its future largely reliant on the sustainability of *ex situ* and reintroduced populations, we undertook an evaluation of the genetic diversity of the global addax population to provide crucial information for the management of the species across the management spectrum (Hoffmann et al., 2015). Biodiversity in the Sahelo-Saharan region is under-studied (Durant et al., 2014), and this evaluation of the genetic diversity of one of its most iconic species adds considerable knowledge to the area. We aim to provide the first global picture of addax genetics, including samples from wild, *ex situ* populations (Europe, North America and UAE), and the reintroduced Tunisian population for use in conservation management decision making.

## Materials and Methods

### Sampling

Faecal samples were collected from wild addax during field surveys in the Tin Toumma region, Niger (2012 - 2017), and Chad (2001). Samples from *ex situ* addax populations, including hair, tissue, and bloods (EDTA), were collected opportunistically during routine veterinary procedures from EEP and SSP institutions in Europe and North America, respectively, and Al Ain Zoo and the Environment Agency - Abu Dhabi (EAD) in the United Arab Emirates. Samples were collected from 104 of 108 censused addax across the three reintroduced populations in Tunisia using biopsy darts. Further sampling details are shown in Table 1 and detailed in Supp. File S1.

**Table 1.** Samples from global addax populations included in this study. Population sizes frequently change, and estimates are provided as a guide only. NA indicates the samples were of insufficient quality and were not included in nuclear DNA analyses.

| Sample origin | Population code | Sample collection dates | Population size estimate at time of sampling | Sample type | Number samples | Number successful at mtDNA | Number successful at nuclear DNA |
| --- | --- | --- | --- | --- | --- | --- | --- |
| Tunisia | Tunisia |  | 108 |  | 104 | 102 | 95 |
| Haddej | Haddej | 2017 | 59 | Tissue | 55 | 55 | 51 |
| Jbil NP | JNP | 2017 | 16 | Tissue | 16 | 15 | 14 |
| Senghar-Jabbes NP | SJNP | 2017 | 33 | Tissue | 33 | 32 | 30 |
| UAE |  |  |  |  | 137 | 125 | 105 |
| Al Ain Zoo | AAZ | 2010-2019 | ~280 | Blood | 67 | 62 | 39 |
| Environment Agency – Abu Dhabi | EAD | 2018-2019 | ~75 | Blood | 70 | 63 | 66 |
| Europe (EEP) |  |  | ~230 |  | 65 | 56 | 34<sup>a</sup> |
| 8 EAZA institutions | EEP | 2012-2019 |  | Hair | 19 | 18 | NA |
|  |  |  |  | Blood/Tissue | 44 | 36 | 34 |
|  |  |  |  | GenBank | 2 | 2 | NA |
| USA (SSP) |  |  | ~225 |  | 43 | 43 | 42<sup>b</sup> |
| 7 AZA institutions | SSP | 2010-2018 |  | Blood/Tissue | 43 | 43 | 42 |
| Wild |  |  | < 100 |  | 37<sup>d</sup> | 32 | 0 |
| Niger (Termit region) | Niger | 2012-2017 |  | Faeces<sup>c</sup> | 34 | 29 | NA |
| Chad (Egeui, now extinct) | Chad | 2010 |  | Faeces<sup>c</sup> | 3 | 3 | NA |
| Total |  |  |  |  | 386 | 358 | 276 |

<sup>a</sup> from 3 institutions
<sup>b</sup> from 6 institutions
<sup>c</sup> Molecular species identification has been carried out and all samples confirmed as addax.
<sup>d</sup> Each sample is assumed to originate from a different individual ( $n = 37$ ). Samples with different mitochondrial lineages cannot originate from the same individual and so an absolute minimum of 11 individuals are present but more likely a minimum of 16 individuals were sampled after accounting for sampling timeframes.

### Mitochondrial DNA analyses

All faecal samples were molecularly identified as addax through sequencing of cytochrome b following Verma and Singh (2002). A 723 bp fragment of the control region was amplified from all sample types using custom designed primers AdnewF (5’–GCTATAGCCCCACTATCAAC) and AdnewR (5’– GCGGGTTGCTGGTTTCACGC). PCR products were sequenced in both directions on an ABI 3130XL genetic analyser - see Supp. File S1 for additional details.

The resulting sequences were aligned along with 11 published control region sequences for addax (GenBank accessions: JN632591 (Hassanin et al. 2012), MZ474955-MZ475965 (Hempel et al. 2021)). Unique haplotypes were identified using the R package HAPLOTYPES (Aktas, 2020).

Alphabetical nomenclature was assigned to haplotypes in contemporary samples, and existing GenBank accession numbers used for haplotypes unique to museum samples. A 76 bp indel was detected in some haplotypes, and the impact of indel re-coding methods were assessed (see Supp. File S1), before selecting the simple index coding method (Simmons & Ochoterena, 2000). The R package PEGAS (Paradis, 2010) was used to estimate nucleotide and haplotype diversity, deviation from neutrality using Tajima’s D with significance estimated under a beta distribution (Paradis, 2020) with Bonferroni correction, and statistical parsimony (TCS) haplotype networks (Templeton et al., 1992).

### Nuclear DNA analyses

A double digest restriction-site associated DNA (ddRAD) library was prepared from samples submitted as blood or tissue following a modified Peterson et al (2012) protocol (see Supp. File S1). Briefly, DNA was fragmented using the restriction enzymes SphI and SbfI, and a unique pair of 5 or 7 bp barcodes was ligated to each sample. The pooled library was then size selected between 400 and 700 bp using gel electrophoretic extraction, prior to PCR amplification to incorporate adapter sequences. Samples were sequenced across eight libraries, and positive controls were used to confirm repeatability within and between libraries. Each library was sequenced (150 bp, paired-end) on a full lane of an Illumina HiSeq.

STACKS v2.52 (Rochette et al., 2019) was used to demultiplex the raw data. Reads were aligned using BWA (Li & Durbin, 2009) to the reference genome of the closely related scimitar-horned oryx (*Oryx dammah* assembly version 1.1, GCF_014754425.2; Humble et al. 2020) and SNPs were called using STACKS within a custom SNAKEMAKE pipeline (Köster & Rahmann, 2012). SNPs on autosomal chromosomes with a minimum genotyping rate of 95% and linkage disequilibrium (LD) R^2^ < 0.5 were retained – known as the relaxed LD SNP set. A stringent LD SNP set was generated for analyses requiring minimum LD (ADMIXTURE) for which only SNPs with R^2^ < 0.2 were retained (1073 SNPs) - see Supp. File S1for additional filtering details.

Population structure was initially assessed using principal component analysis performed within the R package ADEGENET (Jombart, 2008). The package ADMIXTURE (Alexander et al., 2009) was then used with the stringent LD SNP set, assessing *K* 2 to 8 with a 10-fold cross validation with 200 bootstrap resampling runs to estimate the standard errors. Lastly, STRUCTURE was used to provide a Bayesian estimation of population structure (Pritchard et al., 2000), implemented using the package PARALLELSTRUCTURE (Besnier & Glover, 2013). As STRUCTURE assigns clusters by maximising HWE, which is frequently violated in managed populations, and related individuals exist in the dataset which enhances deviation from HWE due to shared haplotypes, we ran STRUCTURE using the relaxed LD SNP set as well as the stringent LD SNP set. The admixture model was run using a burn-in of 50,000 followed by 100,000 repetitions for each of *K* 2 - 10 with 10 repetitions at each value of *K* (with all other parameters as default. Results for both ADMIXTURE and STRUCTURE were visualised using the POPHELPER package for R (Francis, 2017) and CLUMPP (Jakobsson & Rosenberg, 2007). We also performed the ADMIXTURE and STRUCTURE analyses without the Tunisian metapopulation to assess whether ancestry estimation was affected by the large number of samples from this divergent reintroduced population.

Observed heterozygosity (H_O_), expected heterozygosity within populations (H_S_) and the fixation index (F_IS_) were calculated using the HIERFSTAT package (Goudet, 2005). Standardised multilocus heterozygosity (sMLH) was calculated using the INBREEDR package (Stoffel et al., 2016) to generate an estimate of inbreeding which is unaffected by allele frequencies. In addition, F and Fhat3, which are affected by allele frequency estimates, were calculated using PLINK (Chang et al., 2015), and overall levels of relatedness within the population was assessed using the KING-robust kinship estimates calculated in KING (Manichaikul et al., 2010). Allelic richness (Ar) and private allelic richness (pAr) were calculated using ADZE (Szpiech et al., 2008), standardised to N = 14 (the smallest sample size of Jbil NP). Pairwise F_ST_ was calculated using the DARTR package for R (Gruber et al., 2018) implementing the Weir and Cockerham measure (Weir & Cockerham, 1984), with 1000 bootstraps used to estimate 95% confidence intervals. AMOVA was calculated using the R package POPPR (Kamvar et al., 2014) and significance was tested using the randomisation procedure implemented within the R package ADE4 (Dray & Dufour, 2007) using 1000 repeats. All analyses were carried out within R v4.0.3 (R Core Team, 2020).

## Results

### Mitochondrial DNA

A 723 bp fragment of the mitochondrial control region was obtained from 359 samples, and we identified a total of 25 haplotypes (Supp. Table S1, Supp. Fig. S1), 18 identified within contemporary samples. We identified 9 haplotypes within the 29 faecal samples collected from Niger between 2012 and 2017, and two additional haplotypes from three faecal samples collected from Chad in 2001 (Supp. Table S1), resulting in a minimum of 11 individuals represented within these samples. We identified 8 haplotypes from 324 individuals and an additional two GenBank sequences across *ex situ* populations and the reintroduced population in Tunisia. Only haplotype N was detected in the wild and a contemporary population (Tunisia; Supp. Table S1). Of the eight haplotypes identified by Hempel et al (2021) from museum specimens, a single haplotype (M) was also detected in two samples from the presumed extinct population in Chad. Mean haplotype diversity across all haplotypes was 0.799 and mean nucleotide diversity was 0.0047, and diversity estimates are higher for wild populations than *ex situ* and reintroduced.

**Table 3.**
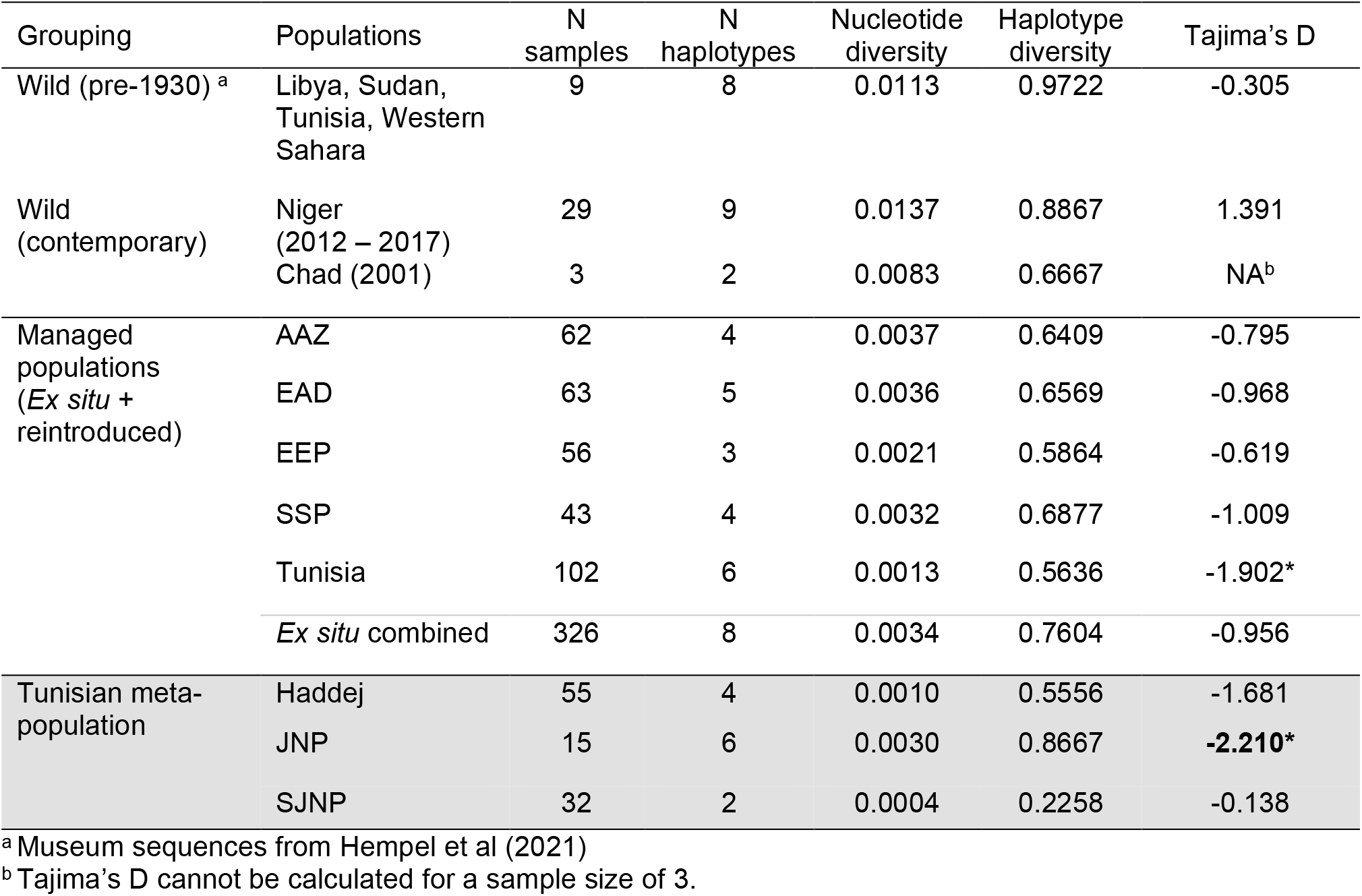
mtDNA diversity between wild and managed populations at 723 bp of d-loop. Significance of Tajima’s D is indicated by * and significance after Bonferroni correction is indicated by bold.

Evolutionary relationships between the observed haplotypes are shown in Figure 2. The 8 haplotypes from *ex situ* and reintroduced populations form a tight central cluster, with haplotype Y being the most divergent (five mutational steps from nearest haplotype). Haplotypes from wild populations were much more diverse. Average nucleotide diversity was also greater in wild haplotypes (pi = 0.014) than *ex situ*/reintroduced haplotypes (pi=0.003).

**Figure 2.**
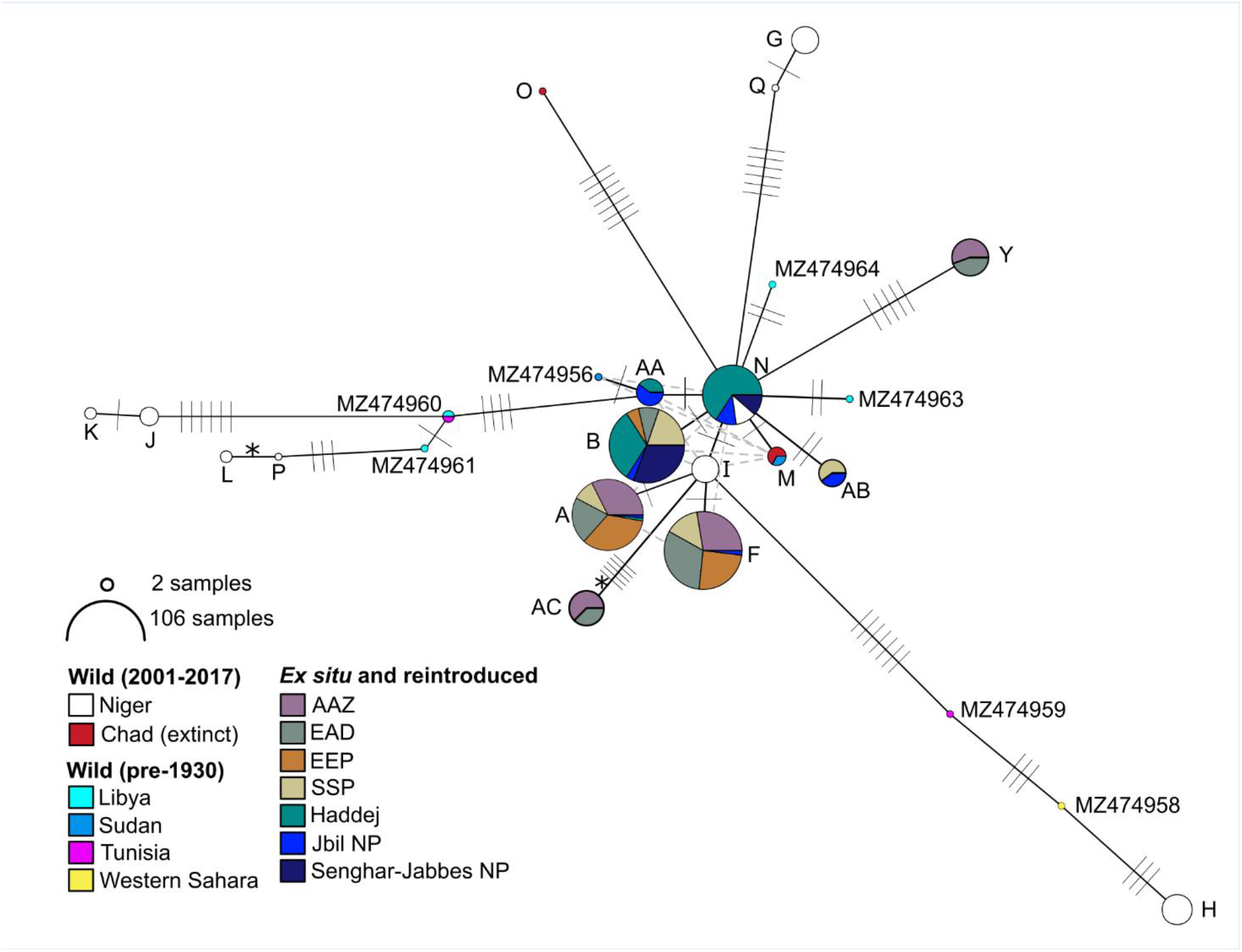
Network of addax mtDNA control region haplotypes. Circles represent haplotypes, with most likely evolutionary relationships indicated by solid black lines, and alternative likely relationships shown by dashed grey lines. Mutational steps are shown by hashes. The 76bp indel was recoded as a single base pair 5^th^ state and is indicated by *, all other indels were excluded. Haplotype circles are coloured according to populations as shown, with circle size indicating sample size (log scale) as indicated. Haplotype nomenclature follows Supp. Table S1. Wild (pre-1930) haplotypes are from Hempel et al (2021).

### Nuclear DNA

We analysed genome-wide nuclear SNP markers for 276 individuals. Across eight ddRAD libraries, 322 samples were sequenced representing 276 individuals, including 18 samples sequenced between 2 and 6 times each as positive controls. On average, 6,226,520 reads were sequenced per sample (range: 14,475 – 29,460,298), with an average of 5,751,874 (91.2%) reads that were mapped against the scimitar-horned oryx genome (range: 39.6% - 97.1%). We identified a total of 24,395 SNPs, and after applying quality filters, the relaxed SNP set retained 1704 SNPs with a genotyping rate of 95% across individuals.

### Global population structure

AMOVA showed that over 83.9% of the variation is partitioned within individuals (Phi = 0.161, *p* = 0.001, Supp. Table S2). There was evidence of variation at the population level, with 12.9% of the variation distributed amongst the five primary *ex situ* populations (Phi = 0.129, *p* = 0.001). F_ST_ ranged between 0.053 (AAZ - EAD) to 0.183 (EAD - Tunisia), shown in Figure 3A, with all measures significant (*p* < 0.001). F_ST_ estimates within Tunisia (Figure 3B) show that divergence is greatest between Jbil NP and Senghar-Jabbes NP (F_ST_ = 0.085 ± 0.009 (95% CI)). F_ST_ estimates within Tunisia were lower than comparisons between the Tunisian metapopulation as a whole and other global populations, though pairwise comparisons including Jbil NP (Jbil NP – Haddej F_ST_ = 0.053; Jbil NP – Senghar-Jabbes NP F_ST_ = 0.085) were similar or greater than estimates between some global populations (Figure 4a, AAZ – EAD F_ST_ = 0.053).

**Figure 3.**
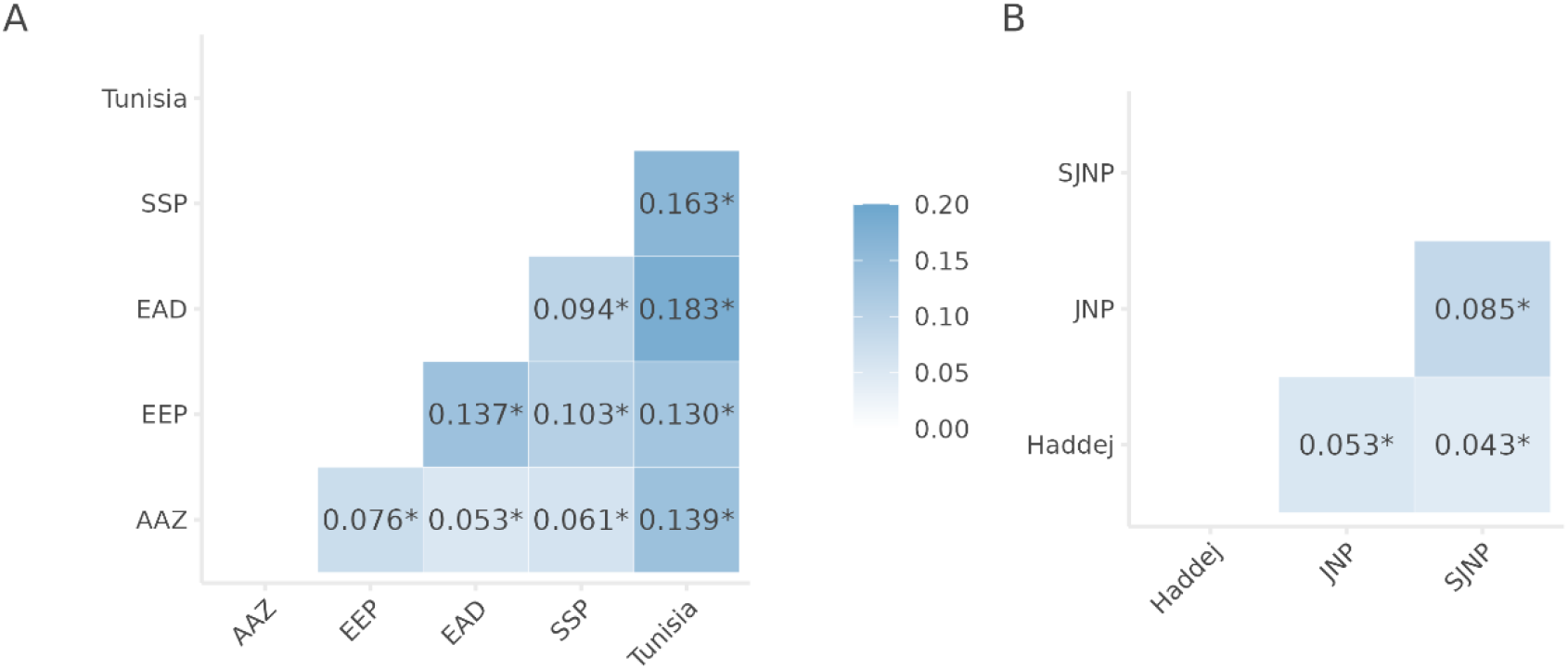
Pairwise F_ST_ estimates for the five primary ex situ populations (a) and for the three Tunisian populations (b). All pairwise F_ST_ estimates were significant (as indicated by *).

**Figure 4.**
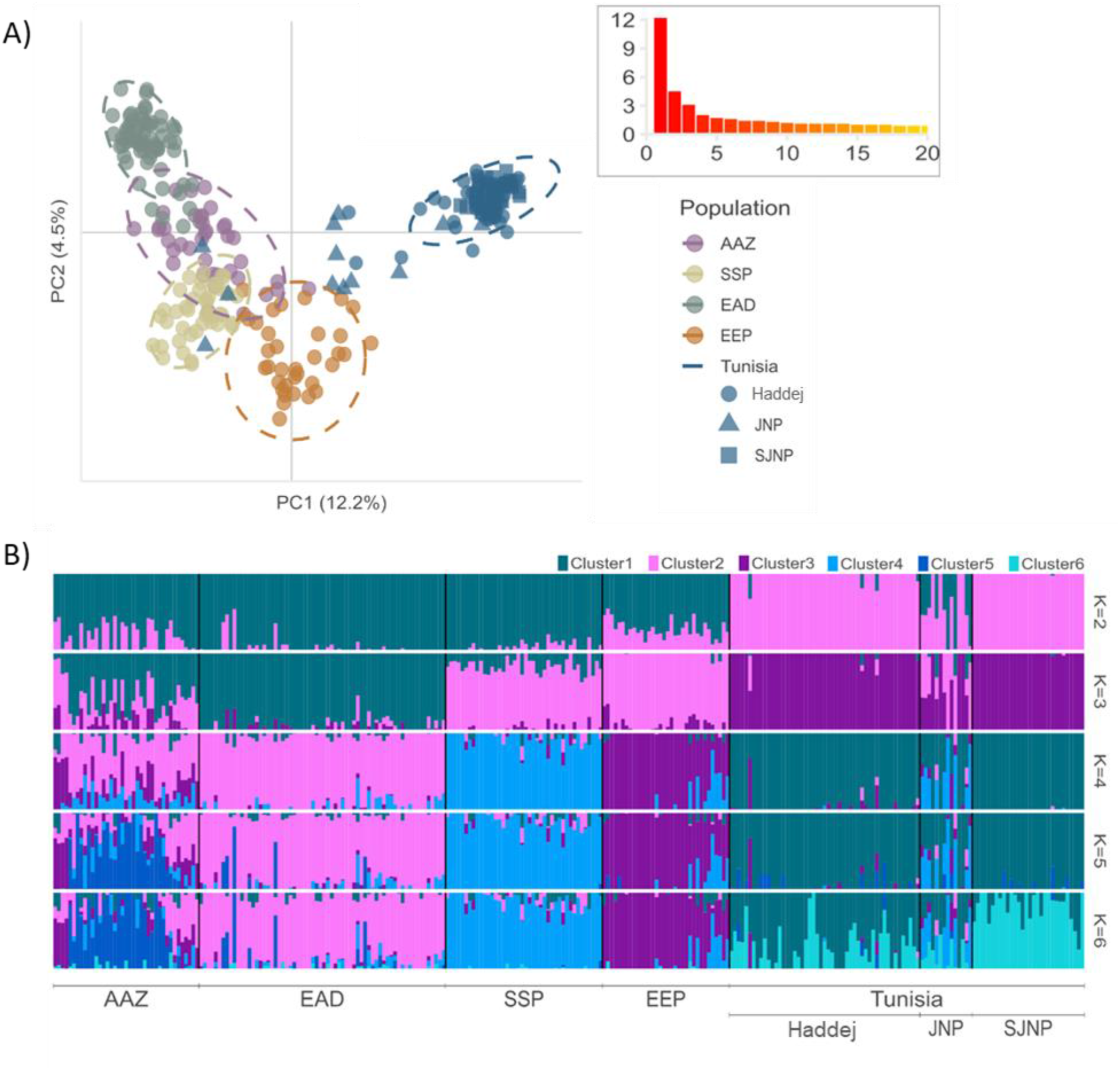
Visualisation of population structure in the ex situ and reintroduced addax populations using two methods. A) PCA using 1704 SNPs in the managed addax populations, showing PC1 and PC2 (percentage of variation explained shown in brackets). The Tunisian metapopulations are represented by different shapes, as shown in the legend. Inset shows the first 20 eigenvalues. B) ADMIXTURE results using 1073 SNPs filtered to minimise linkage disequilibrium, showing, for each individual, the proportion of genetic membership to each ancestry cluster for the most informative number of clusters (K2 to K6).

We carried out an initial assessment of population structure using PCA, shown in Figure 4A. Principal component (PC) 1 accounted for the greatest proportion of variation within the dataset (11.3 %), which primarily separates the Tunisian metapopulation from other global populations. The remaining four *ex situ* populations formed continuums along PC1 and PC2, with none forming independent cluster. This pattern was supported by subsequent PCs (Supp. Fig. S2).

To explore the population structure at a finer scale, we used ADMIXTURE to investigate individual ancestry using 1073 SNPs filtered to minimise linkage disequilibrium. Cross-validation suggested that 5 clusters was the most supported model (Supp. Fig. S3), and results of K=2 to K=6 are shown in Figure 4B. At all estimates of K, Tunisia remained a distinct cluster. The EEP and SSP largely formed an independent cluster at K=3 and become distinct from one another at K=4, although the EEP population included several individuals more similar to the SSP population than to other EEP addax. The two Arabian populations (AAZ and EAD) show ancestry similarly differentiated from other populations, but with varying levels of admixture at all estimates of K. The AAZ is composed of at least two clusters with varying levels of admixture from K=4 onwards. After excluding the Tunisian population, there was no substantial difference in the resulting population structure (Supp. Fig. S4). Additional analyses using STRUCTURE showed broadly similar results (Supp. Fig. S5 and S6) with higher levels of admixture depending upon the degree of filtering to minimise linkage disequilibrium.

Within Tunisia, Jbil NP was found by PCA to include several individuals that did not cluster closely with addax from Haddej or Senghar-Jabbes NP, and three individuals clustering with the SSP (Figure 4A), one of which was a surviving founder translocated from the SSP to Jbil NP in 2007. ADMIXTURE analyses showed evidence of high levels of admixture within the Jbil NP addax (Figure 4B), with genetic signatures from the SSP, the EEP, and the wider Tunisian population at all values of K.

### Genetic diversity within populations

Estimates of genetic diversity were similar across the EAD, EEP and SSP populations, but deviations from Hardy-Weinberg expectations were detected in both AAZ and Tunisia (Figure 5, Supp. Supp. Table S3). For AAZ, observed heterozygosity was significantly lower than expected heterozygosity (H_O_ = 0.168, H_S_ = 0.2), and F_IS_ was raised (F_IS_ = 0.115 ± 0.011 95% CI). The population mean sMLH was in line with the other *ex situ* populations despite high variation in individual sMLH estimates, and pairwise kinship (KING-robust) estimates were very low (mean KING = −0.2). Combined, these diversity estimates are indicative of a Wahlund effect within AAZ.

**Figure 5.**
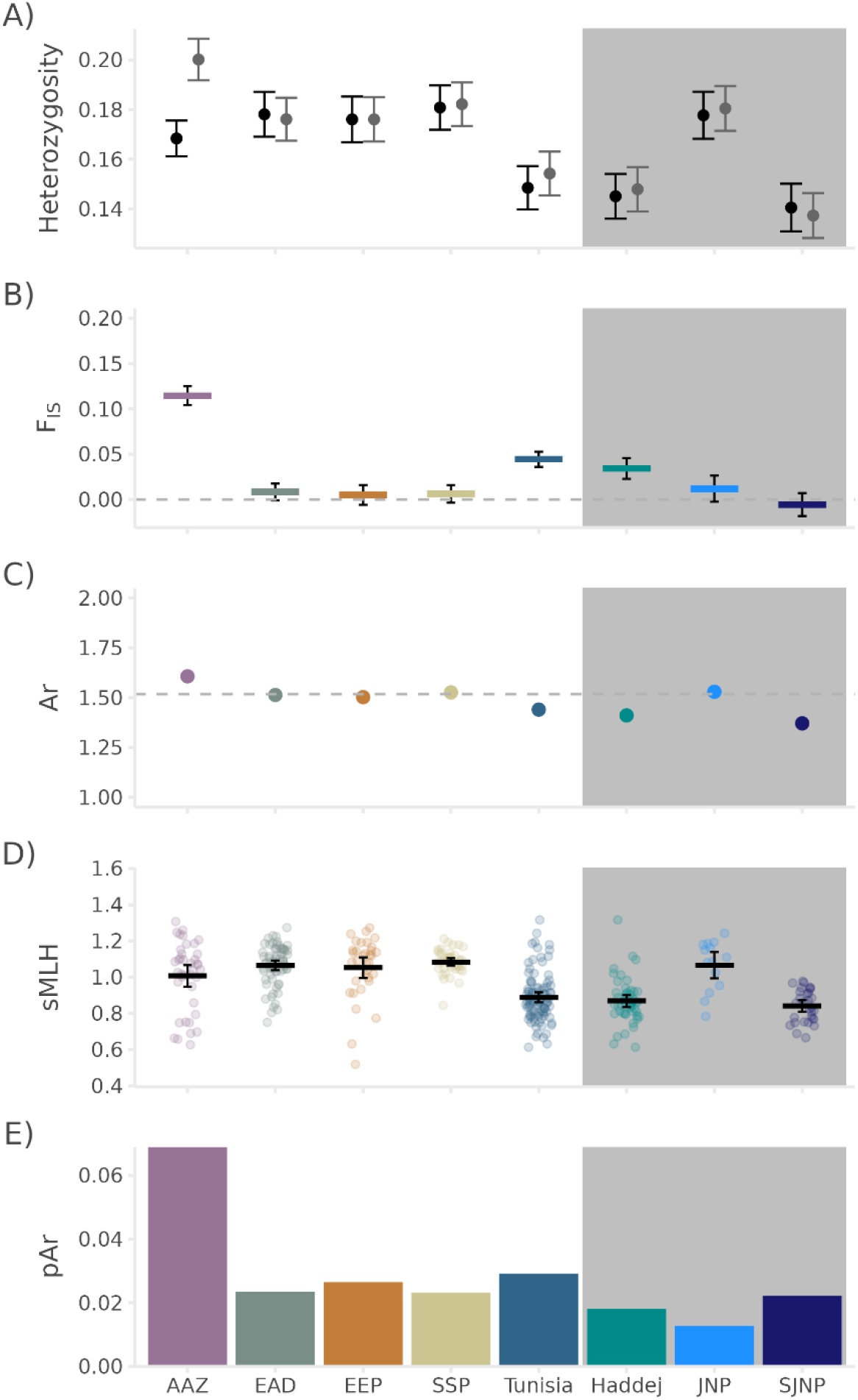
Genetic diversity measures in the ex situ and reintroduced populations. A) Mean observed heterozygosity (black points) and expected heterozygosity (grey points) for each population, with 95% confidence limits (black and grey bars respectively). B) Mean F_IS_ (coloured bars) within each population and 95% confidence limits (black error bars). C) Mean allelic richness (Ar) (points) standardized to N = 14, with 95% confidence limits (black error bars), with the global average indicated by the horizontal dashed line. D) Individual standardised mulitlocus heterozygosity (sMLH). E) Private allelic richness (pAr) standardized to N = 14. In all cases, estimates for Tunisia summarise all three national parks, which are also shown independently (shaded in grey).

Within Tunisia, only F_IS_ was raised (F_IS_ = 0.044 ± 0.008 95% CI) and this was largely driven by Haddej (Figure 5B, Supp. Table S3). sMLH (Figure 3C) was also reduced in Tunisia as a whole, which seems to be driven by the lower individual diversity in Haddej and SJNP compared to JNP. Estimates of inbreeding using F and Fhat3 follow a similar pattern (Supp. Fig. S7A-B), and low levels of kinship (Supp. Fig. S7C) suggests that this pattern is not driven by an excess of related individuals within our dataset.

## Discussion

The data presented here provide a genetic baseline that is needed to underpin sound international population management of addax across the management spectrum from *ex situ* populations that are managed to varying extents, to the remnant population in the wild. The resolution of genetic data is intrinsically tied to sample quality, which in turn is often tightly linked to the type of source population. Challenging field conditions meant that faecal samples were the only sample type available from remnant wild populations, including the now extinct population in Chad and the remnant population in Niger. The mitochondrial data generated from these samples provide a first insight into the genetic status of the remnant wild population, as well as a vital overview between wild, reintroduced, and managed populations. Wild faecal DNA quality was insufficient for ddRAD analysis, but SNP data generated from the higher quality blood and tissue samples collected from managed populations provide data to guide population management decisions at an institutional, regional and a global level. We draw the following conclusions from the analysed dataset:

### In situ genetic diversity is higher than ex situ genetic diversity, but both populations contain unique genetic variants

We detected 11 mtDNA haplotypes from 37 faecal samples collected in the wild, nine of which were detected from the last remaining wild population in Niger over the last 10 years. The two haplotypes detected in three samples from Chad provided an indication of genetic diversity that has been lost in recent years. The dwindling Niger population retains high genetic diversity (nucleotide diversity 0.014 and haplotype diversity 0.887) compared to *ex situ* populations, and although it was not possible to assess levels of inbreeding and genetic load, nor declines in fitness, unique genetic diversity clearly persists within this wild population that has not been captured within the *ex situ* population.

mtDNA diversity in *ex situ* populations was substantially lower than the wild population. We detected only eight haplotypes from 327 samples across 18 institutions over four continents, (nucleotide diversity = 0.0034, haplotype diversity = 0.7604). Only a single haplotype (N) was shared amongst wild and *ex situ* addax. These levels of diversity within the insurance population are substantially lower than closely related arid land Hippotragini species which went Extinct in the Wild, the scimitar-horned oryx (*Oryx dammah*) with 43 control region haplotypes (Ogden et al., 2020) and Arabian oryx (*Oryx leucoryx*) with 11 control region haplotypes (Ochoa et al., 2016). Although the genetic diversity retained within *ex situ* populations is relatively low, potentially limiting its adaptive potential (Franklin, 1980; Spielman et al., 2004), it does include diversity that has been lost from the wild.

Previously analysed mitogenomes revealed weak phylogeographic structure across the historic range of addax (Hempel et al., 2021), and, after incorporating additional contemporary samples from both the wild and captive populations, our analyses of the control region continue to support this lack of geographic signature (Figure 2). Unsampled haplotypes, such as those that are now extinct, impact the accuracy of haplotype network reconstruction (Paradis, 2018) and we struggled to reconstruct evolutionary relationships amongst haplotypes carrying the 76bp indel on haplotypes L and AC. Nevertheless, the networks generated from mitogenomes (Hempel et al., 2021) and the single control region locus analysed, here, were largely congruent. Whilst the TCS network reveals uncertainty surrounding precise placement of several haplotypes from *ex situ* populations (Figure 2), they remain tightly clustered. Mitochondrial diversity within *ex situ* populations, therefore, exhibits lower evolutionary divergence than that which remains in the wild, and likely represents a relatively small component of historical diversity.

Of the eight mtDNA haplotypes detected in *ex situ* populations, no single population contained all haplotypes, indicating population structure. There was a strong degree of overlap between the SSP (four haplotypes) and the EEP (three haplotypes, all detected within SSP). Tunisia’s reintroduced population included all four haplotypes detected in its EEP and SSP source populations, but also included two additional haplotypes (haplotypes AA and N). The absence of haplotypes AA and N from the EEP and SSP within our dataset may be due to under sampling or because they have been lost from these populations. Management of the EEP and SSP populations generally focuses on transferring males among herds of breeding females, which could lead to population structure at this maternally inherited marker resulting in additional diversity at unsampled institutions. Alternatively, if the haplotypes occurred at low frequency within the EEP or SSP at the time, they may have been removed during the translocation to Tunisia or have subsequently died out. This highlights a need for careful and well documented selection of individuals for reintroduction to prevent loss of genetic diversity within source populations that remain as insurance populations.

### Contemporary population structure provides insights into genetic relationships amongst ex situ populations

Population structure was detected amongst the *ex situ* and reintroduced populations of addax, with distinct clusters corresponding to the five primary populations. At four clusters (K=4), addax populations were divided into regional clusters (EEP, SSP, Tunisia and UAE), and as the number of clusters increased from four to six, additional population structure was revealed within and between the two UAE institutions (AAZ and EAD). The structure across all other populations remained consistent after excluding Tunisia from the analysis and largely similar independent of SNP filtering, which is known to affect population structure analyses for ddRAD data with low SNP numbers (B. R. Wright et al., 2019).

Maximising genetic diversity is a priority when selecting founders for reintroductions from available source populations (IUCN/SSC, 2013), a goal which can only be achieved through understanding the genetic relationships between source populations. The founders of the large, coordinated addax breeding programmes, the SSP and EEP, are poorly known (Spevak et al., 1993), and records are lacking for the UAE populations, though some founders are suspected to have originated from the USA private sector (unsampled here but likely genetically similar to the SSP, Hauser et al 2022).

Combining the evidence from the PCA, ADMIXTURE, and diversity analyses, our results support a close genetic relationship amongst all the analysed populations, and exchanges between the SSP and EEP (Enright, 2019) alongside mixing of individuals from these two populations in Tunisia, makes it unlikely that outbreeding would be problematic when mixing populations (Frankham et al., 2011).

Nevertheless, each population retained a distinct signature and novel genetic variation, hinting towards novel founders within the UAE, with high allelic and private allelic richness within AAZ (though these metrics may be inflated due to incomplete admixture, Brockett et al. 2022) and novel, evolutionarily distinct mtDNA haplotypes at AAZ and EAD.

Levels of genetic diversity within *ex situ* and reintroduced populations are relatively low, with limited genomic diversity and structure. Therefore, maximising diversity within a reintroduction by selecting founders from multiple source populations is advised. This also highlights the vital need for integrated management of captive and reintroduced populations to promote geneflow and minimise any further loss of genetic diversity from the insurance populations (Ralls et al., 2018). Managing the partitioned genetic diversity found within *ex situ* addax populations to preserve genetic variants and minimise impacts of localised inbreeding must now be a key objective of future metapopulation planning of the species. Managing populations at an international scale is challenging, particularly for species such as the addax which have both limited founder records and poorly-known pedigrees (Ivy et al., 2016). However, integration of molecular data into population modelling approaches is becoming increasingly feasible (for example see (Gooley et al., 2022; McLennan et al., 2018; Belinda R. Wright et al., 2021) and may be valuable in identifying both optimal and feasible management strategies.

### Semi-managed populations in Tunisia provide a window on the genetic outcomes of reintroduction

The success of reintroduction attempts is reliant on a multitude of factors, including genetic diversity, but are rarely sufficiently assessed over the long term (Brockett et al., 2022; Pierson et al., 2016). Despite being entirely descended from EEP and SSP individuals approximately 6 – 7 generations ago, the Tunisian metapopulation was the most genetically distinct population in our analysis. This was evident at all scales assessed, including two haplotypes undetected in *ex situ* populations (N and AA), four out of the five highest pairwise F_ST_ values (0.130 – 0.183), low heterozygosity estimates (indicative of inbreeding, which can strongly affect F_ST_ estimates, Jost 2008), and consistently formed a unique cluster in structural assessments. The Tunisian metapopulation was founded with only 14 individuals from 2 institutions with a high relatedness amongst founders (Supp. Fig. S8), supporting the genetic evidence of a bottleneck effect at the founding of this population with rapid genetic drift over the 6-7 generations since the founders were translocated to Tunisia in the 1980s. The divergence between the Tunisian metapopulation and its source populations is therefore likely a combination of a founder effect and genetic drift.

Reduction in diversity and heterozygosity is predicted to be more pronounced in smaller populations (Frankham, 1996) and reserves (e.g. Heller et al. 2010), and is not uncommon following founder events, and particularly serial and stepping-stone founder events (Excoffier et al., 2009; Slatkin & Excoffier, 2012) like the formation of the addax metapopulation in Tunisia. Disentangling the bottleneck/founder effects from genetic drift and selective processes is challenging (Belinda R. Wright et al., 2021), particularly in the absence of samples from the founders which would enhance our understanding of how genetic diversity has changed within the metapopulation over time. However, as this study includes a snapshot of almost all addax alive in Tunisia in 2017 and, due to the nature of the translocations with Haddej and Senghar-Jabbes NP only receiving addax descended from the 1980s founders and Jbil NP being the only population to receive a combination of the original and additional founders in 2007, the contemporary partitioning of diversity throughout the metapopulation provides an indication of the success of each strategy. Following the augmentation in 2007 (approximately 1-3 generations ago) using 13 genetically individuals selected from the EEP and SSP following pedigree analyses, Jbil NP was found to have substantially greater mitochondrial and nuclear diversity than its counter parts in Haddej and Senghar-Jabbes NP, despite its smaller population size (Table 1). Notably, Senghar-Jabbes NP had the lowest estimates of population level diversity, and was founded with only 10 individuals from the original Bou Hedma NP/Haddej, supporting the expectations of reduced diversity following a stepping-stone founder event (Excoffier et al., 2009; Slatkin & Excoffier, 2012).

Despite the limited number of genetic founders to Haddej and Senghar-Jabbes NP, these populations have persisted and expanded under management. Species and populations have recovered from the brink of extinction with very small numbers of founders (e.g. Mauritius kestrel (Groombridge et al., 2012), Chatham Island robin (Kennedy et al., 2014) and the black footed ferret (Santymire et al., 2014)), but substantial reductions in genetic diversity are inevitable. Severe bottlenecks and associated reductions in genetic diversity have been widely shown to have long lasting effects on fitness and population persistence (Bouzat, 2010; Frankham, 2005; Ralls et al., 2020). Therefore, when additional genetic variation is available for reintroductions, it is advisable to provide the populations with maximal adaptive potential (IUCN/SSC, 2013). Given the perilous state of wild populations of Sahelo-Saharan megafauna, conservation translocations are an increasingly critical tool in securing populations within their indigenous range. The results from our analyses demonstrate the need to genetically monitor populations post-release and, if appropriate, genetically augment through multiple releases to increase diversity and counteract drift (Dlugosch & Parker, 2008).

Ultimately, the ability of a population to adapt to its environment over the long-term is determined by the amount of diversity in the form of allelic variation (Allendorf, 1986; Caballero & García-Dorado, 2013). Both the mtDNA and nuclear data presented here indicate that the Tunisian meta-population is genetically depauperate compared to the *ex situ* population as a whole and to individual *ex situ* populations. Although the Tunisian populations have remained managed to date, the ultimate aim is to establish a wild, self-sustaining population within the historic range in southern Tunisia. Population viability modelling incorporating the genetic data developed here would be valuable to assess the optimal management strategies to maximise population growth and genetic diversity, and to determine release numbers. Nevertheless, this case study has illustrated the need for both careful selection of founders for reintroduction and ongoing genetic monitoring, and it further illustrates that collection of genetic samples from the outset should be a critical part of any translocation (Brockett et al., 2022), ideally with subsequent submission to a biobank for long-term preservation.

## Conclusions

This work has demonstrated the critical importance of the remaining wild population of addax within Niger as a genetic reservoir as, despite the diminishing size of this population, it retains notably greater genetic diversity than the substantially larger global *ex situ* and reintroduced populations combined. Whilst addax *ex situ* insurance populations have and will undoubtedly continue to play a pivotal role in preventing extinction, conservation efforts must focus on protecting the last remnant population.

At the same time, conservation translocations have proven successful for addax in North Africa and Chad, and genetic management of viable source populations will be critical for long-term reintroduction success. Re-established populations, particularly those that are small and geographically isolated, may require metapopulation management with translocations between sub-populations to minimise any further loss of genetic diversity and maximise long-term population viability. Given that genetic diversity is partitioned within global *ex situ* populations it is urgent that *ex situ* holders cooperate in a global conservation management plan to ensure maximal retention of genetic diversity. Integrated management under a ‘One Plan Approach’ (Traylor-Holzer et al., 2019) is increasingly being advocated for and the Conservation Centers for Species Survival (C2S2) has prioritised addax as a species for using genomics to connect private and zoological populations (Wildt et al., 2019). Decisions regarding optimal strategies at both regional and international scales will benefit from improved integration of molecular genetic data, as developed here, into population viability modelling, and group management strategies may be valuable for integrating populations with varying intensities of management (Gooley et al., 2022). Particularly, this study provides the necessary data to model conservation translocation outcomes in Tunisia and inform management decisions.

There is a clear need to improve genetic resources for addax to facilitate conservation management, and biobanking, cell line generation and reproductive technologies must be prioritised for this species. Improved availability of high-quality genetic samples from key insurance populations is vital for developing the data to enable integrated population management. Given the global nature of these insurance populations and challenges associated with international translocations, movement of germ cells may prove crucial to establishing effective geneflow. Such resources may prove invaluable to the conservation of addax in the long term, but, nevertheless, the need for *in-situ* conservation action and improved global population management of *ex situ* populations is immediate.

## Acknowledgements

We thank all zoological institutions that provided samples, Fabian Krause from Zoo Hannover for his assistance in sourcing samples from the EEP and Asako Navarro from San Diego Zoo Wildlife Alliance for assistance with DNA samples from SSP institutions. We thank Jennifer Kaden and Muhammad Ghazali for technical laboratory analyses. We are grateful to the Direction Générale des Forêts (Ministère de l’Agriculture, des Ressources Hydrauliques et de la Pêche Maritime) for permission to collect tissue samples from addax using biopsy darting in Tunisia, to the CRDA of Tataouine, Kebili and Gafsa and to the staff of Senghar-Jabbes, Jbil and BouHedma/Haddej NP for their assistance. We thank, Habib Kammoun, Nadia Chelbi, Youssef Derouiche, and Mohsen Ben Nejma. We are grateful to Director General H.E Ghanim Alhajeri, Al Ain Zoo, for providing funds and for supporting this work. We thank Issaka Houdou and Alkabouss Matchano (SaharaConservation) and the late Ali Abba Gana (Projet Gestion Durable de la Biodiversité et des Aires Protégées, Ministère de l’Environnement et de la Lutte Contre la Désertification, Niger) for assistance with field missions in Niger during which samples were collected. We thank Rachel Gardner at Marwell Wildlife for producing maps of addax historical and contemporary ranges. We thank David Mallon for reviewing and improving the manuscript.

Funding for this research was provided by the institutions listed in author affiliations. Al Ain Zoo, the Environment Agency - Abu Dhabi, Marwell Wildlife (UK registered charity number 275433), Royal Zoological Society of Scotland (UK registered charity number SC004064), SaharaConservation, San Diego Zoo Wildlife Alliance. However, the funders did not have any additional role in study design, data collection and analysis, decision to publish, or preparation of the manuscript.

## Data availability

Mitochondrial control region haplotypes were submitted to GenBank under accession numbers XXX – XXX. The raw, demultiplexed ddRAD data for this study have been deposited in the European Nucleotide Archive (ENA) at EMBL-EBI under accession number PRJEB53821. Analysis code, including the custom SNAKEMAKE pipeline for SNP calling, are available on GitHub (https://github.com/XXX) and will be uploaded to FigShare/Dryad upon acceptance.

## Literature cited

Aktas, C. (2020). haplotypes: Manipulating DNA Sequences and Estimating Unambiguous Haplotype Network with Statistical Parsimony (R package version 1.1.2). https://cran.r-project.org/package=haplotypes

Alexander, D. H., Novembre, J., & Lange, K. (2009). Fast model-based estimation of ancestry in unrelated individuals. Genome Research, 19(9), 1655–1664. https://doi.org/10.1101/gr.094052.109

Allendorf, F. W. (1986). Genetic drift and the loss of alleles versus heterozygosity. Zoo Biology, 5(2), 181–190. https://doi.org/10.1002/ZOO.1430050212

Armstrong, E., Leizagoyen, C., Martínez, A. M., González, S., Delgado, J. V., & Postiglioni, A. (2011). Genetic structure analysis of a highly inbred captive population of the African antelope Addax nasomaculatus. Conservation and management implications. Zoo Biology, 30(4), 399–411. https://doi.org/10.1002/zoo.20341

Ballou, J. D., & Lacy, R. C. (1995). Identifying genetically important individuals for management of genetic variation in pedigreed populations. In J. D. Ballou, M. Gilpin, & T. J. Foose (Eds.), Population Management for Survival and Recovery (pp. 76–111). Columbia University Press.

Bertram, B. C. R. (1988). Re-introducing scimitar-horned oryx into Tunisia. In A. Dixon & D. Jones (Eds.), Conservation and Biology of Desert Antelope (pp. 136–145). Christopher Helm.

Besnier, F., & Glover, K. A. (2013). ParallelStructure: A R Package to Distribute Parallel Runs of the Population Genetics Program STRUCTURE on Multi-Core Computers. PLoS ONE, 8(7), e70651. https://doi.org/10.1371/journal.pone.0070651

Bouzat, J. L. (2010). Conservation genetics of population bottlenecks: The role of chance, selection, and history. Conservation Genetics, 11(2), 463–478. https://doi.org/10.1007/S10592-010-0049-0

Brito, J. C., Godinho, R., Martínez-Freiría, F., Pleguezuelos, J. M., Rebelo, H., Santos, X., Vale, C. G., Velo-Antón, G., Boratyński, Z., Carvalho, S. B., Ferreira, S., Gonçalves, D. V., Silva, T. L., Tarroso, P., Campos, J. C., Leite, J. V., Nogueira, J., Álvares, F., Sillero, N., … Carranza, S. (2014). Unravelling biodiversity, evolution and threats to conservation in the Sahara-Sahel. Biological Reviews, 89(1), 215–231. https://doi.org/10.1111/brv.12049

Brockett, B., Banks, S., Neaves, L. E., Gordon, I. J., Pierson, J. C., & Manning, A. D. (2022). Establishment, persistence and the importance of longitudinal monitoring in multi-source reintroductions. Animal Conservation. https://doi.org/10.1111/acv.12764

Caballero, A., & García-Dorado, A. (2013). Allelic diversity and its implications for the rate of adaptation. Genetics, 195(4), 1373–1384. https://doi.org/10.1534/genetics.113.158410

Chang, C. C., Chow, C. C., Tellier, L. C., Vattikuti, S., Purcell, S. M., & Lee, J. J. (2015). Second-generation PLINK: rising to the challenge of larger and richer datasets. GigaScience, 4(1), 7. https://doi.org/10.1186/s13742-015-0047-8

Chardonnet, P. (2007). Translocation of addax and oryx: a big step towards the restoration of Saharan wildlife in Tunisia. Gnusletter, 26, 13.

Convention on Biological Diversity. (2020). Updated zero draft of the post-2020 global biodiversity framework.

Correll, T., & Houston, W. (1999). Addax. Gnusletter, 18, 56.

Crnokrak, P., & Roff, D. A. (1999). Inbreeding depression in the wild. Heredity, 83(3), 260–270. https://doi.org/10.1038/sj.hdy.6885530

Dlugosch, K. M., & Parker, I. M. (2008). Founding events in species invasions: genetic variation, adaptive evolution, and the role of multiple introductions. Molecular Ecology, 17, 431–449. https://doi.org/10.1111/J.1365-294X.2007.03538.X

Dray, S., & Dufour, A. B. (2007). The ade4 package: Implementing the duality diagram for ecologists. Journal of Statistical Software, 22(4), 1–20. https://doi.org/10.18637/jss.v022.i04

Durant, S. M., Wacher, T., Bashir, S., Woodroffe, R., De Ornellas, P., Ransom, C., Newby, J., Abáigar, T., Abdelgadir, M., El Alqamy, H., Baillie, J., Beddiaf, M., Belbachir, F., Belbachir-Bazi, A., Berbash, A. A., Bemadjim, N. E., Beudels-Jamar, R., Boitani, L., Breitenmoser, C., … Pettorelli, N. (2014). Fiddling in biodiversity hotspots while deserts burn? Collapse of the Sahara’s megafauna. Diversity and Distributions, 20(1), 114–122. https://doi.org/10.1111/ddi.12157

Enright, W. (2019). International Addax Studbook.

Excoffier, L., Foll, M., & Petit, R. J. (2009). Genetic Consequences of Range Expansions. Annual Review of Ecology, Evolution, and Systematics, 40, 481–501. https://doi.org/10.1146/ANNUREV.ECOLSYS.39.110707.173414

Francis, R. M. (2017). pophelper: an R package and web app to analyse and visualize population structure. Molecular Ecology Resources, 17(1), 27–32. https://doi.org/10.1111/1755-0998.12509

Frankham, R. (1995). Conservation genetics. Annual Review of Genetics, 29(1), 305–327.

Frankham, R. (1996). Relationship of Genetic Variation to Population Size in Wildlife. Conservation Biology, 10(6), 1500–1508. https://doi.org/10.1046/J.1523-1739.1996.10061500.X

Frankham, R. (2005). Genetics and extinction. In Biological Conservation (Vol. 126, Issue 2, pp. 131– 140). https://doi.org/10.1016/j.biocon.2005.05.002

Frankham, R., Ballou, J. D., & Briscoe, D. A. (2010). Introduction to Conservation Genetics (Second). Cambridge University Press.

Frankham, R., Ballou, J. D., Eldridge, M. D. B., Lacy, R. C., Ralls, K., Dudash, M. R., & Fenster, C. B. (2011). Predicting the Probability of Outbreeding Depression. Conservation Biology, 25(3), 465– 475. https://doi.org/10.1111/J.1523-1739.2011.01662.X

Franklin, I. R. (1980). Evolutionary change in small populations. In M. E. Soulé & B. A. Wilcox (Eds.), Conservation Biology: an evolutionary-ecological perspective. (pp. 135–149). Sinauer Associates.

Gilbert, T., Woodfine, T., Petretto, M., Nouioui, M., Houston, B., & Riordan, P. (2018). Reintroduction of addax to Djebil National Park, Tunisia. In P. S. Soorae (Ed.), Global Reintroduction Perspectives: 2018. Case studies from around the globe (pp. 120–124). IUCN/SSC Reintroduction Specialist Group, Gland, Switzerland and Environment Agency, Abu Dhabi, UAE. https://doi.org/https://doi.org/10.2305/IUCN.CH.2018.08.en

Gooley, R. M., Dicks, K. L., Ferrie, G. M., Lacy, R. C., Ballou, J. D., Callicrate, T., Senn, H., Koepfli, K.-P., Edwards, C. W., & Pukazhenthi, B. S. (2022). Applying genomics to metapopulation management in North American insurance populations of southern sable antelope (Hippotragus niger niger) and addra gazelle (Nanger dama ruficollis). Global Ecology and Conservation, 33, e01969. https://doi.org/10.1016/J.GECCO.2021.E01969

Gooley, R. M., Tamazian, G., Castañeda-Rico, S., Murphy, K. R., Dobrynin, P., Ferrie, G. M., Haefele, H., Maldonado, J. E., Wildt, D. E., Pukazhenthi, B. S., Edwards, C. W., & Koepfli, K.-P. (2020). Comparison of genomic diversity and structure of sable antelope (Hippotragus niger) in zoos, conservation centers, and private ranches in North America. Evolutionary Applications, 13(8), 2143–2154. https://doi.org/10.1111/eva.12976

Goudet, J. (2005). HIERFSTAT, a package for R to compute and test hierarchical F-statistics. Molecular Ecology Notes, 5(1), 184–186. https://doi.org/10.1111/j.1471-8286.2004.00828.x

Groombridge, J. J., Raisin, C., Bristol, R., & Richardson, D. S. (2012). Genetic Consequences of Reintroductions and Insights from Population History. In J. G. Ewen, D. P. Armstrong, K. A. Parker, & P. J. Seddon (Eds.), Reintroduction Biology: Integrating Science and Management (pp. 395–440). Blackwell Publishing. https://doi.org/10.1002/9781444355833.ch12

Gruber, B., Unmack, P. J., Berry, O. F., & Georges, A. (2018). dartr: An r package to facilitate analysis of SNP data generated from reduced representation genome sequencing. Molecular Ecology Resources, 18(3), 691–699. https://doi.org/10.1111/1755-0998.12745

Heller, R., Okello, J. B. A., & Siegismund, H. (2010). Can small wildlife conservancies maintain genetically stable populations of large mammals? Evidence for increased genetic drift in geographically restricted populations of Cape buffalo in East Africa. Molecular Ecology, 19(7), 1324–1334. https://doi.org/10.1111/J.1365-294X.2010.04589.X

Hempel, E., Westbury, M. V., Grau, J. H., Trinks, A., Paijmans, J. L. A., Kliver, S., Barlow, A., Mayer, F., Müller, J., Chen, L., Koepfli, K.-P., Hofreiter, M., & Bibi, F. (2021). Diversity and Paleodemography of the Addax (Addax nasomaculatus), a Saharan Antelope on the Verge of Extinction. Genes 2021, Vol. 12, Page 1236, 12(8), 1236. https://doi.org/10.3390/GENES12081236

Hoban, S., Bruford, M., D’Urban Jackson, J., Lopes-Fernandes, M., Heuertz, M., Hohenlohe, P. A., Paz-Vinas, I., Sjögren-Gulve, P., Segelbacher, G., Vernesi, C., Aitken, S., Bertola, L. D., Bloomer, P., Breed, M., Rodríguez-Correa, H., Funk, W. C., Grueber, C. E., Hunter, M. E., Jaffe, R., … Laikre, L. (2020). Genetic diversity targets and indicators in the CBD post-2020 Global Biodiversity Framework must be improved. Biological Conservation, 248, 108654. https://doi.org/10.1016/j.biocon.2020.108654

Hoffmann, A., Griffin, P., Dillon, S., Catullo, R., Rane, R., Byrne, M., Jordan, R., Oakeshott, J., Weeks, A., Joseph, L., Lockhart, P., Borevitz, J., & Sgrò, C. (2015). A framework for incorporating evolutionary genomics into biodiversity conservation and management. Climate Change Responses, 2(1), 1–24. https://doi.org/10.1186/s40665-014-0009-x

Hogg, C. J., Lee, A. V., Srb, C., & Hibbard, C. (2017). Metapopulation management of an Endangered species with limited genetic diversity in the presence of disease: the Tasmanian devil Sarcophilus harrisii. International Zoo Yearbook, 51(1), 137–153. https://doi.org/10.1111/IZY.12144

Humble, E., Dobrynin, P., Senn, H., Chuven, J., Scott, A. F., Mohr, D. W., Dudchenko, O., Omer, A. D., Colaric, Z., Lieberman Aiden, E., Al Dhaheri, S. S., Wildt, D., Oliaji, S., Tamazian, G., Pukazhenthi, B. S., Ogden, R., & Koepfli, K.-P. (2020). Chromosomal-level genome assembly of the scimitar-horned oryx: Insights into diversity and demography of a species extinct in the wild. Molecular Ecology Resources, 00, 1–14. https://doi.org/10.1111/1755-0998.13181

IUCN/SSC. (2013). Guidelines for Reintroductions and Other Conservation Translocations. Version 1.0.

IUCN SSC Antelope Specialist Group. (2016). Addax nasomaculatus. The IUCN Red List of Threatened Species 2016. https://dx.doi.org/10.2305/IUCN.UK.2016-2.RLTS.T512A50180603.en

IUCN SSC Antelope Specialist Group. (2020). IUCN mission to Niger for the conservation of the last wild addax and dama gazelles and the Termit and Tin Toumma National Nature Reserve : Report and Recommendations.

Ivy, J. A., Putnam, A. S., Navarro, A. Y., Gurr, J., & Ryder, O. A. (2016). Applying SNP-derived molecular coancestry estimates to captive breeding programs. Journal of Heredity, 107(5), 403–412. https://doi.org/10.1093/jhered/esw029

Jakobsson, M., & Rosenberg, N. A. (2007). CLUMPP: A cluster matching and permutation program for dealing with label switching and multimodality in analysis of population structure. Bioinformatics, 23(14), 1801–1806. https://doi.org/10.1093/bioinformatics/btm233

Jombart, T. (2008). Adegenet: A R package for the multivariate analysis of genetic markers. Bioinformatics, 24(11), 1403–1405. https://doi.org/10.1093/bioinformatics/btn129

Jost, L. (2008). GST and its relatives do not measure differentiation. Molecular Ecology, 17(18). https://doi.org/10.1111/j.1365-294X.2008.03887.x

Kamvar, Z. N., Tabima, J. F., & Grünwald, N. J. (2014). Poppr: An R package for genetic analysis of populations with clonal, partially clonal, and/or sexual reproduction. PeerJ, 2, e281. https://doi.org/10.7717/peerj.281

Kennedy, E. S., Grueber, C. E., Duncan, R. P., & Jamieson, I. G. (2014). Severe inbreeding depression and no evidence of purging in an extremely inbred wild species—the Chatham Island black robin. Evolution, 68(4), 987–995. https://doi.org/10.1111/EVO.12315

Köster, J., & Rahmann, S. (2012). Snakemake-a scalable bioinformatics workflow engine. Bioinformatics, 28(19), 2520–2522. https://doi.org/10.1093/bioinformatics/bts480

Krause, F. (2016). Addax antelope (Addax nasomaculatus) European studbook.

Lachapelle, J., Colegrave, N., & Bell, G. (2017). The effect of selection history on extinction risk during severe environmental change. Journal of Evolutionary Biology, 30(10), 1872–1883. https://doi.org/10.1111/jeb.13147

Laikre, L., Allendorf, F. W., Aroner, L. C., Baker, C. S., Gregovich, D. P., Hansen, M. M., Jackson, J. A., Kendall, K. C., McKelvey, K., Neel, M. C., Olivieri, I., Ryman, N., Schwartz, M. K., Bull, R. S., Stetz, J. B., Tallmon, D. A., Taylor, B. L., Vojta, C. D., Waller, D. M., & Waples, R. S. (2010). Neglect of genetic diversity in implementation of the convention on biological diversity: Conservation in practice and policy. Conservation Biology, 24, 86–88. https://doi.org/10.1111/j.1523-1739.2009.01425.x

Lande, R. (1994). Risk of population extinction from fixation of new deleterious mutations. Evolution, 48(5), 1460–1469. https://doi.org/10.1111/J.1558-5646.1994.TB02188.X

Lande, Russell, & Shannon, S. (1996). The role of genetic variation in adaptation and population persistence in a changing environment. Evolution, 50(1), 434–437. https://doi.org/10.1111/j.1558-5646.1996.tb04504.x

Li, H., & Durbin, R. (2009). Fast and accurate short read alignment with Burrows-Wheeler transform. Bioinformatics, 25(14), 1754–1760. https://doi.org/10.1093/bioinformatics/btp324

Mallon, D., & Chardonnet, P. (2020). Antelope news: Texotics. Gnusletter, 37(2), 46.

Manichaikul, A., Mychaleckyj, J. C., Rich, S. S., Daly, K., Sale, M., & Chen, W. M. (2010). Robust relationship inference in genome-wide association studies. Bioinformatics, 26(22), 2867–2873. https://doi.org/10.1093/bioinformatics/btq559

Marshall, T. C., Sunnucks, P., Spalton, J. A., Greth, A., & Pemberton, J. M. (1999). Use of genetic data for conservation management: the case of the Arabian oryx. Animal Conservation, 2, 269– 278. https://doi.org/10.1111/j.1469-1795.1999.tb00073.x

McGowan, P. J. K., Traylor-Holzer, K., & Leus, K. (2017). IUCN Guidelines for Determining When and How Ex Situ Management Should Be Used in Species Conservation. Conservation Letters, 10(3), 361–366. https://doi.org/10.1111/CONL.12285

McLennan, E. A., Gooley, R. M., Wise, P., Belov, K., Hogg, C. J., & Grueber, C. E. (2018). Pedigree reconstruction using molecular data reveals an early warning sign of gene diversity loss in an island population of Tasmanian devils (Sarcophilus harrisii). Conservation Genetics, 19(2), 439– 450. https://doi.org/10.1007/S10592-017-1017-8

Müller, H. P., & Engel, H. (2004). The reintroduction of herbivores to Souss Massa National Park, Morocco. In T. Gilbert & T. Woodfine (Eds.), The biology, husbandry and conservation of scimitar-horned oryx (Oryx dammah) (pp. 77–81). Marwell Preservation Trust.

Ochoa, A., Wells, S. A., West, G., Al-Smadi, M., Redondo, S. A., Sexton, S. R., & Culver, M. (2016). Can captive populations function as sources of genetic variation for reintroductions into the wild? A case study of the Arabian oryx from the Phoenix Zoo and the Shaumari Wildlife Reserve, Jordan. Conservation Genetics, 17(5), 1145–1155. https://doi.org/10.1007/s10592-016-0850-5

Ogden, R., Chuven, J., Gilbert, T., Hosking, C., Gharbi, K., Craig, M. S., Al Dhaheri, S. S., & Senn, H. (2020). Benefits and pitfalls of captive conservation genetic management: Evaluating diversity in scimitar-horned oryx to support reintroduction planning. Biological Conservation, 241, 108244. https://doi.org/10.1016/j.biocon.2019.108244

Paradis, E. (2010). Pegas: An R package for population genetics with an integrated-modular approach. In Bioinformatics (Vol. 26, Issue 3, pp. o419–420). https://doi.org/10.1093/bioinformatics/btp696

Paradis, E. (2018). Analysis of haplotype networks: The randomized minimum spanning tree method. Methods in Ecology and Evolution, 9(5), 1308–1317. https://doi.org/10.1111/2041-210X.12969

Paradis, E. (2020). Population Genomics with R. In Population Genomics with R. Chapman and Hall/CRC. https://doi.org/10.1201/9780429466700

Payne, B. L., & Bro-Jørgensen, J. (2016). Disproportionate Climate-Induced Range Loss Forecast for the Most Threatened African Antelopes. Current Biology, 26(9), 1200–1205. https://doi.org/10.1016/j.cub.2016.02.067

Peterson, B. K., Weber, J. N., Kay, E. H., Fisher, H. S., & Hoekstra, H. E. (2012). Double Digest RADseq: An Inexpensive Method for De Novo SNP Discovery and Genotyping in Model and Non-Model Species. PLoS ONE, 7(5), e37135. https://doi.org/10.1371/journal.pone.0037135

Pierson, J. C., Coates, D. J., Oostermeijer, J. G. B., Beissinger, S. R., Bragg, J. G., Sunnucks, P., Schumaker, N. H., & Young, A. G. (2016). Genetic factors in threatened species recovery plans on three continents. Frontiers in Ecology and the Environment, 14(8), 433–440. https://doi.org/10.1002/FEE.1323

Pritchard, J. K., Stephens, M., & Donnelly, P. (2000). Inference of Population Structure Using Multilocus Genotype Data. Genetics, 155(2), 945–959. https://doi.org/10.1534/genetics.116.195164

R Core Team. (2020). R: A language and environment for statistical computing. R Foundation for Statistical Computing. https://www.r-project.org/

Rabeil, T., Garba, H. H. M., Harouna, A., Abagana, A., & Bello, I.. (2016). Addax aerial and ground survey in Niger. 16th Annual Sahelo-Saharan Interest Group Meeting.

Ralls, K., Ballou, J. D., Dudash, M. R., Eldridge, M. D. B., Fenster, C. B., Lacy, R. C., Sunnucks, P., & Frankham, R. (2018). Call for a Paradigm Shift in the Genetic Management of Fragmented Populations. Conservation Letters, 11(2), e12412. https://doi.org/10.1111/conl.12412

Ralls, K., Sunnucks, P., Lacy, R. C., & Frankham, R. (2020). Genetic rescue: A critique of the evidence supports maximizing genetic diversity rather than minimizing the introduction of putatively harmful genetic variation. Biological Conservation, 251(August), 108784. https://doi.org/10.1016/j.biocon.2020.108784

Rochette, N. C., Rivera-Colón, A. G., & Catchen, J. M. (2019). Stacks 2: Analytical methods for paired-end sequencing improve RADseq-based population genomics. Molecular Ecology, 28(21), 4737–4754. https://doi.org/10.1111/mec.15253

Santymire, R. M., Livieri, T. M., Branvold-Faber, H., & Marinari, P. E. (2014). The Black-Footed Ferret: On the Brink of Recovery? In W.. Holt, J. Brown, & P. Comizzoli (Eds.), Reproductive Sciences in Animal Conservation. Advances in Experimental Medicine and Biology (Vol. 753, pp. 119–134). Springer. https://doi.org/10.1007/978-1-4939-0820-2_7

SCF. (2020). Sahara Conservation Fund 2019 Annual Report.

Secretary of the Convention on Biological Diversity. (1992). Convention on Biological Diversity.

Simmons, M. P., & Ochoterena, H. (2000). Gaps as Characters in Sequence-Based Phylogenetic Analyses on JSTOR. Systematic Biology, 49(2), 369–381. https://www.jstor.org/stable/2585224

Slatkin, M., & Excoffier, L. (2012). Serial Founder Effects During Range Expansion: A Spatial Analog of Genetic Drift. Genetics, 191, 171–181. https://doi.org/10.1534/GENETICS.112.139022

Species360. (2022). Zoological Information Management System (ZIMS). https://species360.org

Spevak, E. M., Blumer, E. S., & Correll, T. L. (1993). Species survival plan contributions to research and reintroduction of Addax Addax nasomaculatus. International Zoo Yearbook, 32(1), 91–98. https://doi.org/10.1111/j.1748-1090.1993.tb03520.x

Spielman, D., Brook, B. W., & Frankham, R. (2004). Most species are not driven to extinction before genetic factors impact them. Proceedings of the National Academy of Sciences of the United States of America, 101(42), 15261–15264. https://doi.org/10.1073/pnas.0403809101

Stabach, J. A., Rabeil, T., Turmine, V., Wacher, T., Mueller, T., & Leimgruber, P. (2017). On the brink of extinction-Habitat selection of addax and dorcas gazelle across the Tin Toumma desert, Niger. Diversity and Distributions, 23(6), 581–591. https://doi.org/10.1111/ddi.12563

Stoffel, M. A., Esser, M., Kardos, M., Humble, E., Nichols, H. J., David, P., & Hoffman, J. I. (2016). inbreedR: an R package for the analysis of inbreeding based on genetic markers. Methods in Ecology and Evolution, 7(11), 1331–1339. https://doi.org/10.1111/2041-210X.12588

Szpiech, Z. A., Jakobsson, M., & Rosenberg, N. A. (2008). ADZE: A rarefaction approach for counting alleles private to combinations of populations. Bioinformatics, 24(21), 2498–2504. https://doi.org/10.1093/bioinformatics/btn478

Templeton, A. R., Crandall, K. A., & Sing, C. F. (1992). A cladistic analysis of phenotypic associations with haplotypes inferred from restriction endonuclease mapping and DNA sequence data. III. Cladogram estimation. Genetics, 132(2), 619–633. https://doi.org/10.1093/GENETICS/132.2.619

Traylor-Holzer, K., Leus, K., & Bauman, K. (2019). Integrated Collection Assessment and Planning (ICAP) workshop: Helping zoos move toward the One Plan Approach. Zoo Biology, 38(1), 95– 105. https://doi.org/10.1002/ZOO.21478

Verma, S. K., & Singh, L. (2002). Novel universal primers establish identity of an enormous number of animal species for forensic application. Molecular Ecology Notes, 3(1), 220–222. https://doi.org/10.1046/j.1471-8286

Weir, B. S., & Cockerham, C. C. (1984). Estimating F-statistics for the analysis of population structure. Evolution, 38(6), 1358–1370. https://doi.org/10.1111/j.1558-5646.1984.tb05657.x

Wildt, D., Miller, P., Koepfli, K. P., Pukazhenthi, B., Palfrey, K., Livingston, G., Beetem, D., Shurter, S., Gregory, J., Takács, M., & Snodgrass, K. (2019). Breeding Centers, Private Ranches, and Genomics for Creating Sustainable Wildlife Populations. In BioScience (Vol. 69, Issue 11). https://doi.org/10.1093/biosci/biz091

Willi, Y., & Hoffmann, A. A. (2009). Demographic factors and genetic variation influence population persistence under environmental change. Journal of Evolutionary Biology, 22, 124–133. https://doi.org/10.1111/j.1420-9101.2008.01631.x

Wright, B. R., Grueber, C. E., Lott, M. J., Belov, K., Johnson, R. N., & Hogg, C. J. (2019). Impact of reduced-representation sequencing protocols on detecting population structure in a threatened marsupial. Molecular Biology Reports, 46(5), 5575–5580. https://doi.org/10.1007/S11033-019-04966-6/FIGURES/2

Wright, Belinda R., Hogg, C. J., McLennan, E. A., Belov, K., & Grueber, C. E. (2021). Assessing evolutionary processes over time in a conservation breeding program: a combined approach using molecular data, simulations and pedigree analysis. Biodiversity and Conservation 2021 30:4, 30(4), 1011–1029. https://doi.org/10.1007/S10531-021-02128-4

