## Supplementary material for "Genetic diversity in global populations of the Critically Endangered addax (*Addax nasomaculatus*) and its implications for conservation": Supp. Tables

Dicks et al. *Genetic diversity in the Critically Endangered addax*
Supplementary tables

*Table S1. Number and location of each haplotype detected from samples collected here and sequences available on GenBank. A single additional haplotype AJ235310 exists on GenBank (unpublished) but does not have locational information.*

| | *In situ* (wild) | | *Ex situ* populations | | | | | Tunisia reintroduction | | | Museum samples (1821 – 1926; Hempel et al. 2021) | | | | Total samples | Associated accession number(s) |
|---|---|---|---|---|---|---|---|---|---|---|---|---|---|---|---|---|
| Haplotype | Niger | Chad[a] | AAZ | EAD | EEP | SSP | Tunisia | Haddej | JNP | SJNP | Sudan | Western Sahara | Tunisia | Libya | | |
| A | - | - | 23 | 15 | 24 | 7 | 2 | 1 | 1 | - | - | - | - | - | 71 | |
| B | - | - | - | 8 | 5 | 18 | 60 | 29 | 3 | 28 | - | - | - | - | 91 | |
| F | - | - | 29 | 33 | 27[b] | 15 | 2 | - | 2 | - | - | - | - | - | 106 | MZ474965 / JN632591 |
| G | 5 | - | - | - | - | - | - | - | - | - | - | - | - | - | 5 | |
| H | 6 | - | - | - | - | - | - | - | - | - | - | - | - | - | 6 | |
| I | 5 | - | - | - | - | - | - | - | - | - | - | - | - | - | 5 | |
| J | 3 | - | - | - | - | - | - | - | - | - | - | - | - | - | 3 | |
| K | 2 | - | - | - | - | - | - | - | - | - | - | - | - | - | 2 | |
| L | 2 | - | - | - | - | - | - | - | - | - | - | - | - | - | 2 | |
| M | - | 2 | - | - | - | - | - | - | - | - | 1 | - | - | - | 3 | MZ474955 |
| N | 4 | - | - | - | - | - | 31 | 23 | 4 | 4 | - | - | - | - | 35 | |
| O | - | 1 | - | - | - | - | - | - | - | - | - | - | - | - | 1 | |
| P | 1 | - | - | - | - | - | - | - | - | - | - | - | - | - | 1 | |
| Q | 1 | - | - | - | - | - | - | - | - | - | - | - | - | - | 1 | |
| Y | - | - | 5 | 4 | - | - | - | - | - | - | - | - | - | - | 9 | |
| AA | - | - | - | - | - | - | 5 | 2 | 3 | - | - | - | - | - | 5 | |
| AB | - | - | - | - | - | 3 | 2 | - | 2 | - | - | - | - | - | 5 | |
| AC | - | - | 5 | 3 | - | - | - | - | - | - | - | - | - | - | 8 | |
| MZ474956 | - | - | - | - | - | - | - | - | - | - | 1 | - | - | - | 1 | MZ474956 |
| MZ474958 | - | - | - | - | - | - | - | - | - | - | - | 1 | - | - | 1 | MZ474958 |
| MZ474959 | - | - | - | - | - | - | - | - | - | - | - | - | 1 | - | 1 | MZ474959 |
| MZ474960 | - | - | - | - | - | - | - | - | - | - | - | - | 2 | - | 2 | MZ474960/MZ474962 |
| MZ474961 | - | - | - | - | - | - | - | - | - | - | - | - | - | 1 | 1 | MZ474961 |
| MZ474963 | - | - | - | - | - | - | - | - | - | - | - | - | - | 1 | 1 | MZ474963 |
| MZ474964 | - | - | - | - | - | - | - | - | - | - | - | - | - | 1 | 1 | MZ474964 |
| Total haplotypes | 9 | 2 | 4 | 5 | 3 | 4 | 6 | 4 | 6 | 2 | | | | | | |

[a] Now presumed extinct
[b] Includes GenBank sequences MZ474965 (Hempel et al. 2021) and JN632591 (Hassanin et al. 2012).

*Table S2. Results of the hierarchical analysis of molecular variance (AMOVA), partitioning the variation amongst populations, within individuals within each population, and finally within individuals.*

| Source of variation | df | Sum of squares | % of variation | Phi | *p*-value |
|---|---|---|---|---|---|
| Within samples | 276 | 78128.31 | 83.9 | 0.161 | 0.001 |
| Between samples within populations | 271 | 82597.53 | 3.2 | 0.037 | 0.002 |
| Among populations | 4 | 19596.67 | 12.9 | 0.129 | 0.001 |
| Total | 551 | 180322.51 | 100.0 | NA | NA |

*Table S3. Genetic diversity statistics for the five major ex situ populations of addax, calculated from the 1704 SNPs with a 95% genotyping rate. N is the sample size per population, Ho is the observed heterozygosity, Hs is the expected heterozygosity, and Fis is the fixation index. Allelic richness (Ar) and private allelic richness (Pr) were estimated for sample sizes of 14 (the smallest sample size when the three Tunisian national parks are considered independently) and 34 (the smallest sample size when considering Tunisia as a single population). 95% confidence limits are shown in brackets.*

| Population | N | $H_O$ | $H_S$ | $F_{IS}$ | sMLH | Ar (N=14) | pAr (N=14) |
|---|---|---|---|---|---|---|---|
| AAZ | 39 | 0.168 | 0.200 | 0.115 | 1.007 | 1.607 | 0.069 |
| [95% CL] | | [0.161 - 0.176] | [0.192 - 0.209] | [ 0.104 - 0.125] | [0.947 - 1.067] | [1.6036 - 1.6072] | [0.0687 - 0.0690] |
| EAD | 66 | 0.178 | 0.176 | 0.008 | 1.065 | 1.513 | 0.023 |
| [95% CL] | | [0.169 - 0.187] | [0.167 - 0.185] | [-0.001 - 0.018] | [1.038 - 1.092] | [1.5125 - 1.5135] | [0.0234 - 0. 0236] |
| EEP | 34 | 0.176 | 0.176 | 0.005 | 1.053 | 1.502 | 0.026 |
| [95% CL] | | [0.167 - 0.185] | [0.167 - 0.185] | [-0.006 - 0.016] | [0.996 - 1.110] | [1.5016 - 1.5026] | [0.0264 - 0.0266] |
| SSP | 42 | 0.181 | 0.182 | 0.006 | 1.085 | 1.526 | 0.023 |
| [95% CL] | | [0.172 - 0.190] | [0.173 - 0.191] | [-0.003 - 0.016] | [1.064 - 1.105] | [1.5254 - 1.5264] | [0.0231 - 0.0233] |
| Tunisia | 95 | 0.148 | 0.154 | 0.044 | 0.889 | 1.440 | 0.029 |
| [95% CL] | | [0.140 - 0.157] | [0.145 - 0.163] | [ 0.036 - 0.053] | [0.862 - 0.916] | [1.04393 - 1.4402] | [0.0291 - 0.0293] |
| Haddej | 51 | 0.145 | 0.148 | 0.034 | 0.868 | 1.411 | 0.018 |
| [95% CL] | | [0.136 - 0.154] | [0.139 - 0.157] | [ 0.023 - 0.046] | [0.835 - 0.901] | [1.4101 - 1.4111] | [0.0180 - 0.0182] |
| JNP | 14 | 0.178 | 0.18 | 0.012 | 1.067 | 1.529 | 0.013 |
| [95% CL] | | [0.168 - 0.187] | [0.171 - 0.189] | [-0.002 - 0.026] | [0.994 - 1.139] | [1.5288 - 1.5298] | [0.0127 - 0.0129] |
| SJNP | 30 | 0.14 | 0.137 | -0.006 | 0.841 | 1.371 | 0.022 |
| [95% CL] | | [0.131 - 0.150] | [0.128 - 0.146] | [-0.018 - 0.007] | [0.809 - 0.874] | [1.3702 - 1.3712] | [0.0222 - 0.0223] |
