## Supplementary material for "Genetic diversity in global populations of the Critically Endangered addax (*Addax nasomaculatus*) and its implications for conservation": Supp. Figures

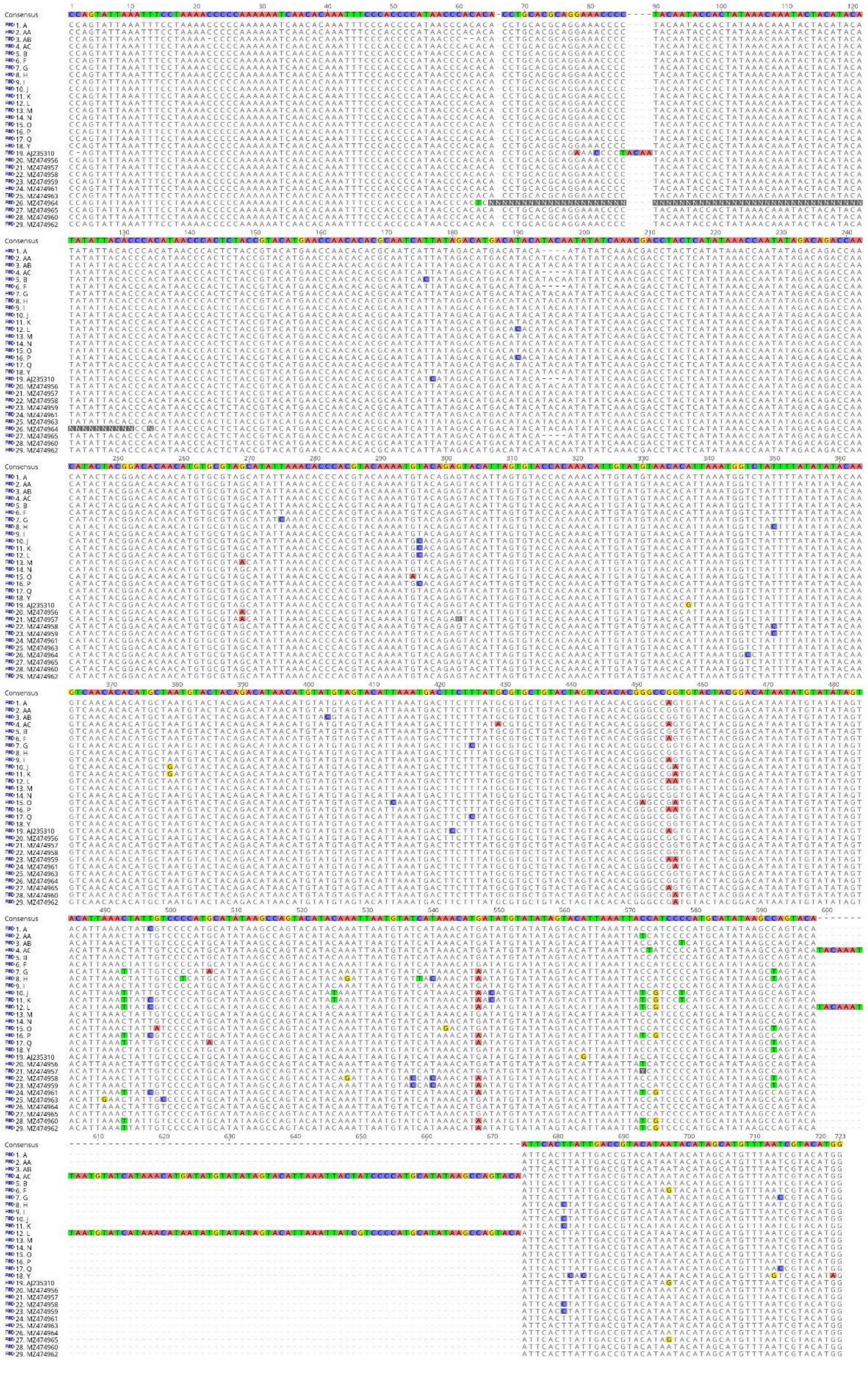

Figure S1. DNA sequence of each haplotype across the 723 bp of the control region analysed. Dashes indicate indels.

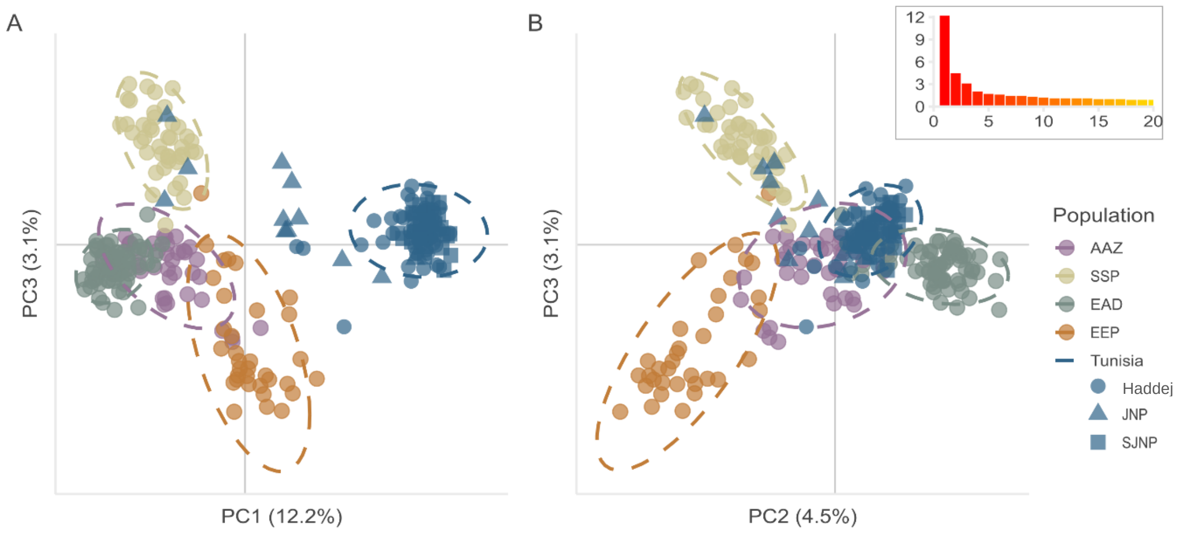


Figure S2. Visualisation of PCA using 1704 SNPs in the managed addax populations, showing A) PC1 and PC3 and B) PC2 and PC3 (percentage of variation explained shown in brackets). The Tunisian metapopulations are represented by different shapes, as shown in the legend. Inset shows the first 20 eigenvalues.

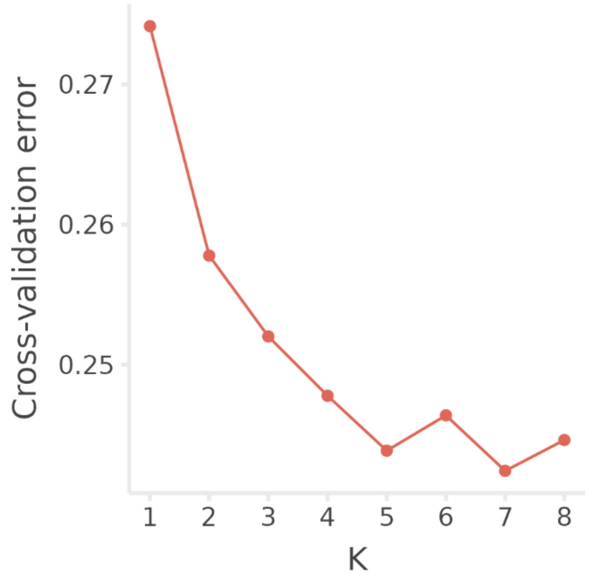


Figure S3. Cross-validation error results for ADMIXTURE. The best supported number of clusters are between K=4 and K=6.

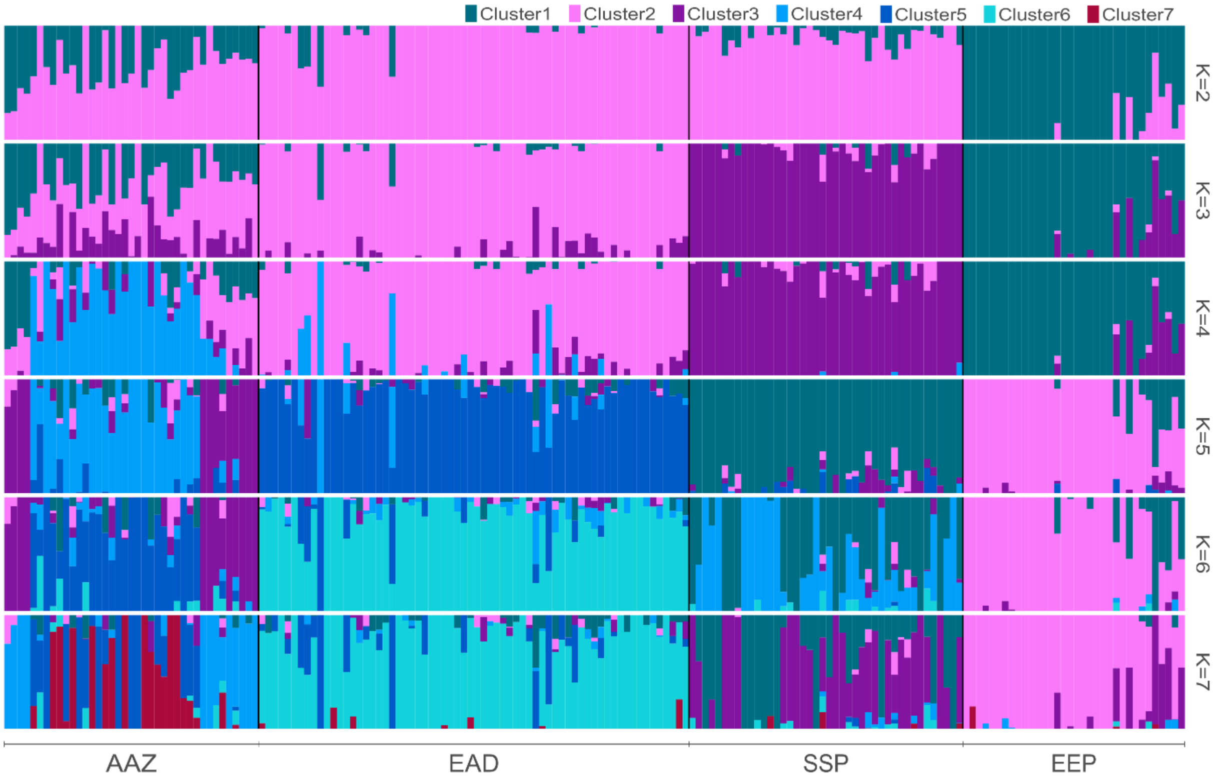


Fig S4. ADMIXTURE results for the four captive populations with Tunisia excluded, using 774 SNPs that were filtered to minimise linkage disequilibrium,

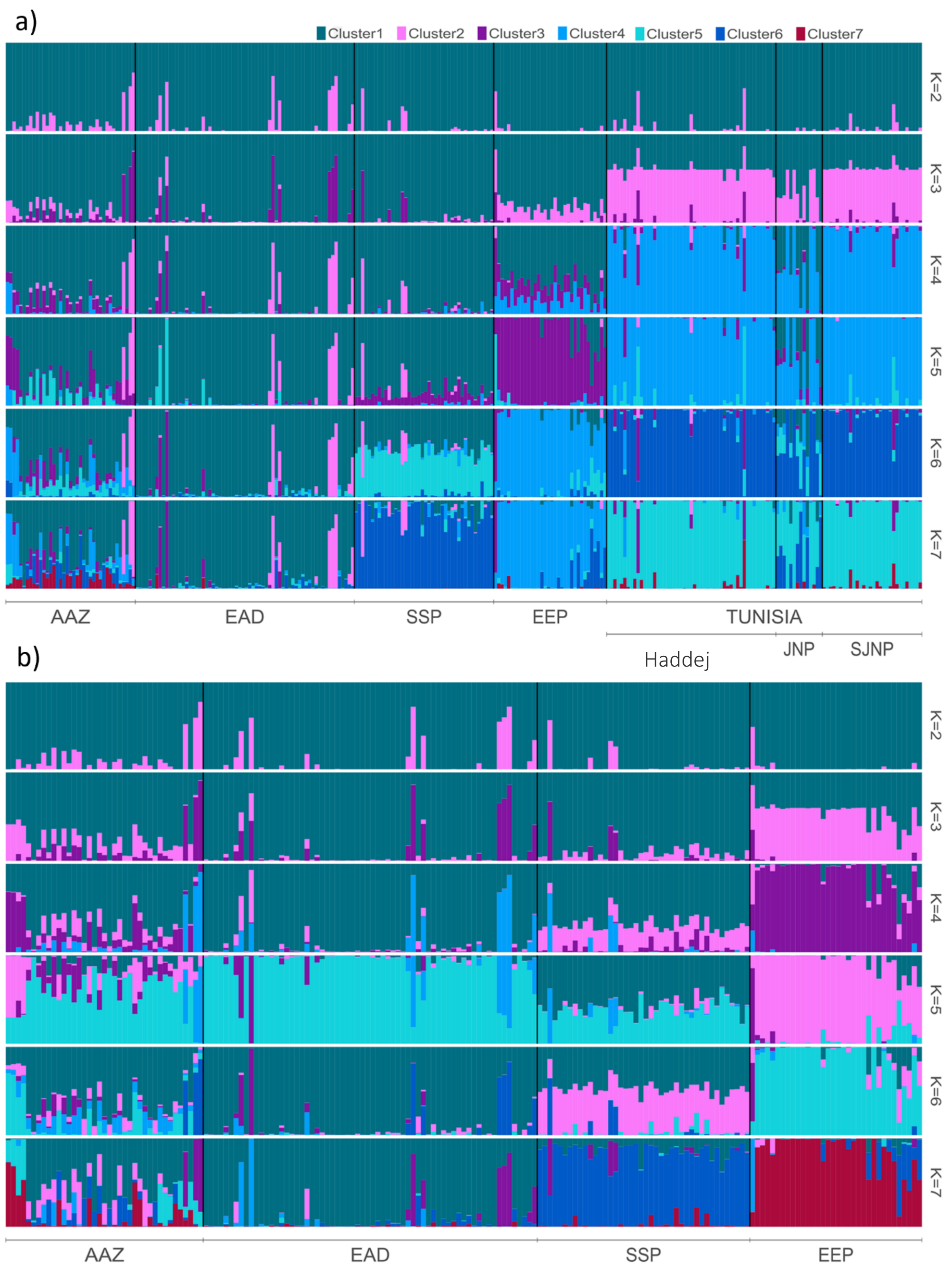


Figure S5. STRUCTURE using the 774 SNPs filtered to minimise linkage disequilibrium for all managed populations (a) and excluding Tunisia (b).

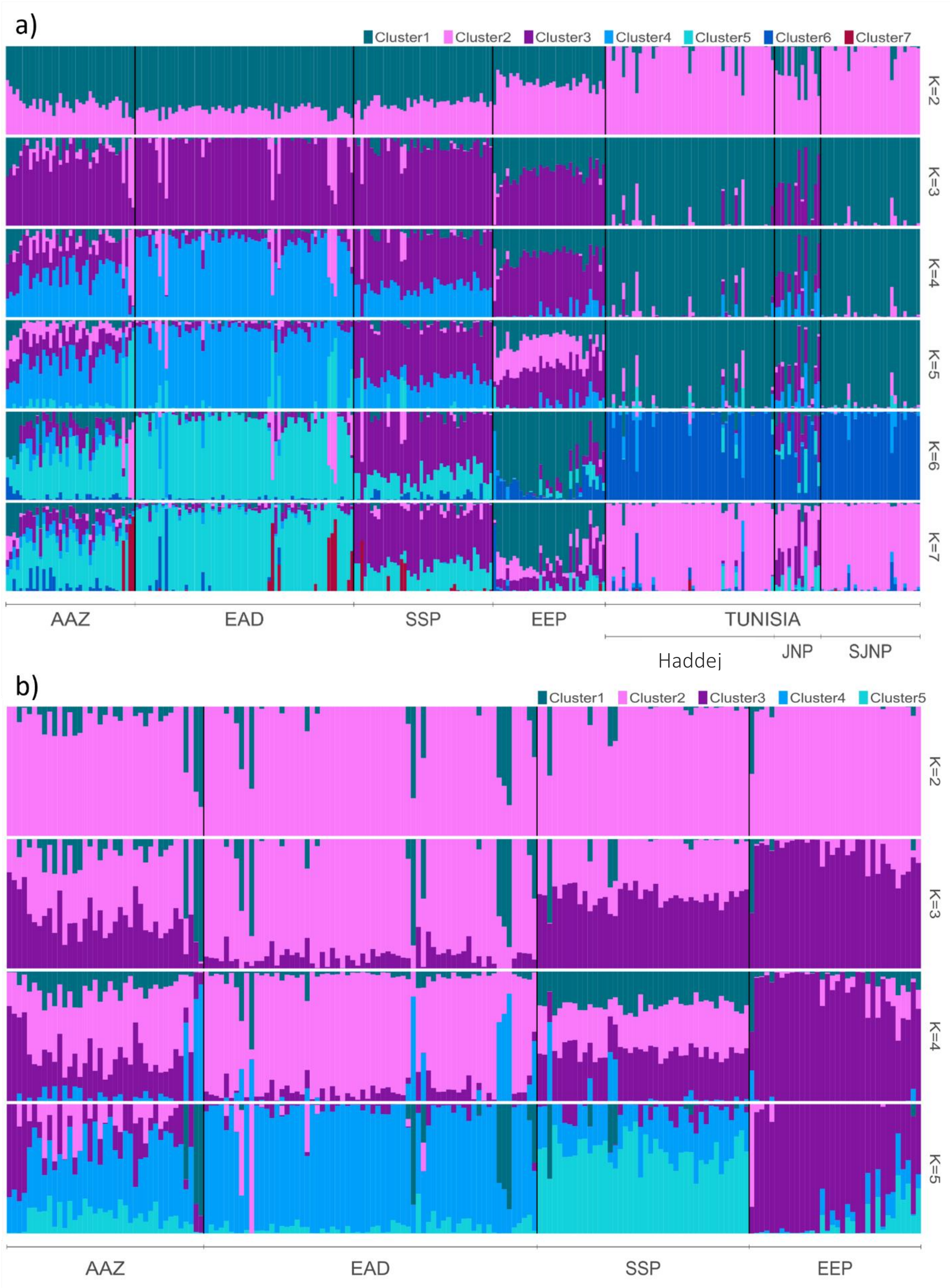


Figure S6. STRUCTURE using all 1704 SNPs (no additional linkage disequilibrium filters applied) for all managed populations (a) and excluding Tunisia (b).

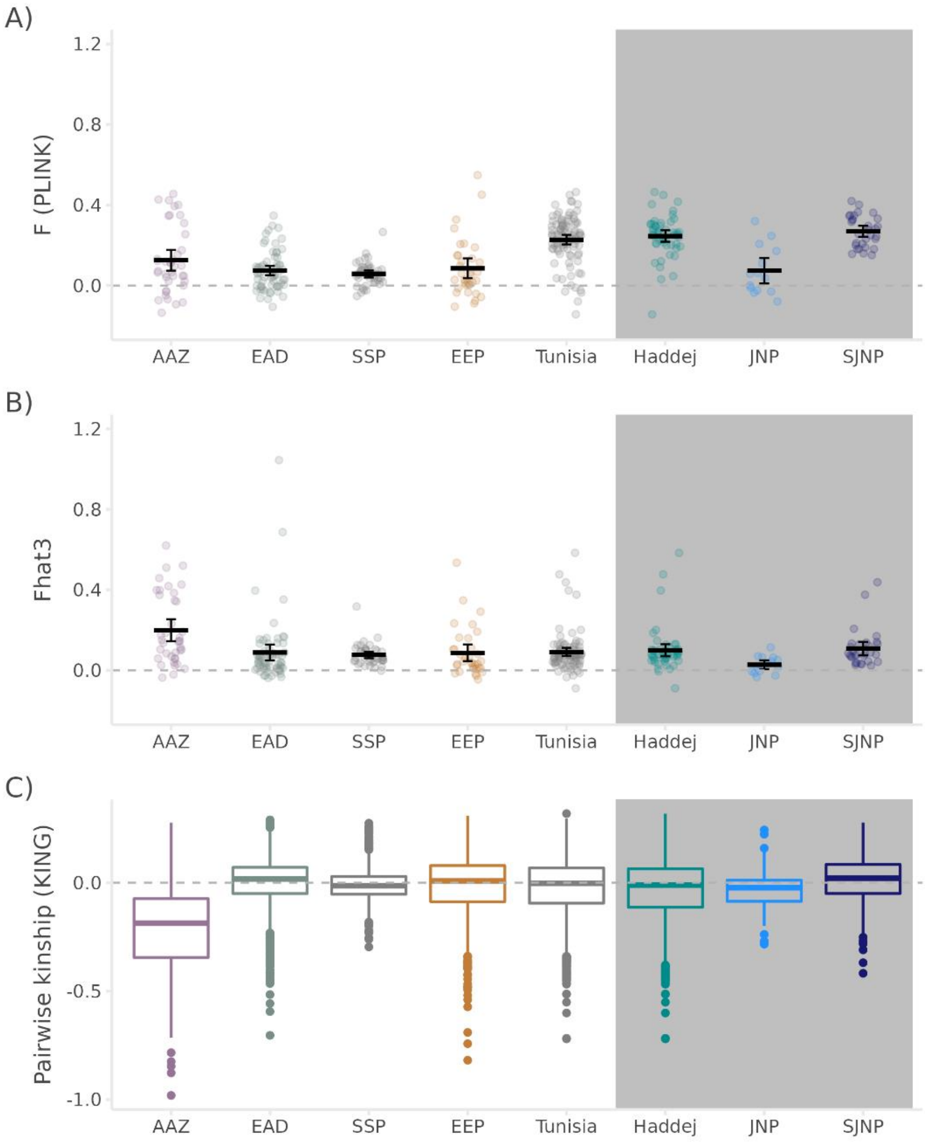


Figure S7. Additional estimates of inbreeding within the managed populations, measured as A) F and B) Fhat3 (using PLINK) with mean estimates are shown as black bars with 95% confidence intervals. C) Boxplots of KING-robust pairwise relatedness, where the centre lines indicate population medians, bounds of the boxes indicate 75% percentiles and whiskers are the range of the data. Estimates of zero (dashed grey line) indicate unrelated individuals for all three measures. Note that KING estimates of relatedness are negatively biased when pairs of individuals are drawn from different populations, which is likely the cause of the increased proportion of negative kinship estimates for AAZ.

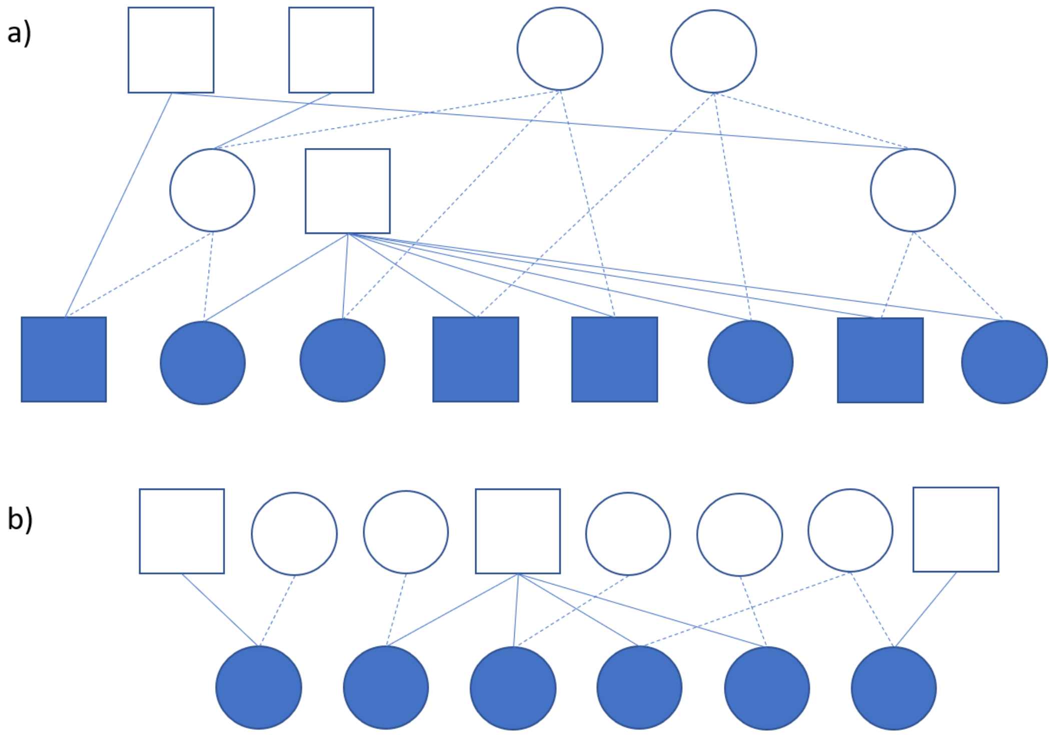


Figure S8. Known pedigree relationships of the 14 addax (dark shapes) used to establish the Tunisian metapopulation in the 1980s from a) the SSP and b) the EEP. First degree relatives (open shapes) are shown, with secondary relationships noted where relevant. Males are represented by squares and females by circles.
