## Supplementary material for "Genetic diversity in global populations of the Critically Endangered addax (*Addax nasomaculatus*) and its implications for conservation": Supp. File S1

### Supplementary methods

#### Sample collection

Samples from *ex situ* addax populations were collected opportunistically during routine veterinary procedures from EEP and SSP institutions in Europe and North America, respectively, and Al Ain Zoo and the Environment Agency - Abu Dhabi (EAD) in the United Arab Emirates. Blood samples were collected in EDTA vacutainers. Tissue samples were stored in 90-100% ethanol either from post-mortem muscle samples or ear punches collected during tagging. Hair samples were stored dry in envelopes.

Tissue samples from the reintroduced Tunisian metapopulation were collected using biopsy darts in the three protected areas in Tunisia over 11 days in 2017 (Jbil National Park – 05 - 06 August and 12-13 December; Senghar-Jabbes National Park – 02-03 October; and Haddej National Park – 09-13 November). Biopsy samples were collected using a Dan-Inject® rifle, model JM Special, using a protocol modified from Karesh et al.(1987). Biopsy darts were equipped with 25mm needles for adult and sub-adult addax, and 20mm needles for juveniles. We individually identified addax using unique facial and horn features together with photographs to minimise unintentional sample duplication. We attempted to sample every adult (>2 years), sub-adult (1-2 years) and juvenile (3 months - 1 year) addax in the population (*N*=110). Calves (<3 months old) and neonates (<1 month old) were not sampled. Biopsy samples were extracted from the needles in the field and preserved in 90% ethanol and held, with the exception of three samples in bent needles that were preserved in the needle in 90% ethanol. Throughout field sampling and storage in Tunisia, all samples were kept at approximately -18°C.

All faecal, blood and tissue samples were stored at -20 °C and hair samples at room temperature upon receipt at RZSS WildGenes prior to laboratory analyses. All samples from outside the UK were imported under the Scottish Government’s Trade in Animals and Related Products (Scotland) Regulations 2012 import permits issued to RZSS. All samples from EAZA institutes were received from European Union institutions between 2012 and 2019 and CITES permits were not required. DNA samples from AZA institutions were loaned to RZSS (CITES Scientific Institution GB039) by San Diego Zoo Global (CITES Scientific Institution US 203). Samples from AAZ and EAD institutions in the UAE were imported under the CITES export and import licenses listed below (note that more than one individual/sample can be listed on a permit):

| Origin Institution/Country | CITES Export Permit : Import Permit |
|---|---|
| Al Ain Zoo (AAZ), United Arab Emirates | 10MEW1621 : 457160/02 |
| | 17MEW7836 : 562290/01 |
| | 17MEW7836 : 562290/02 |
| | 17MEW7836 : 562290/03 |
| | 17MEW7836 : 562290/04 |
| | 17MEW7837 : 562290/05 |
| | 17MEW7837 : 562290/06 |
| | 17MEW7837 : 562290/07 |
| | 17MEW7837 : 562290/08 |
| | 17MEW7838 : 562290/09 |
| | 17MEW7838 : 562290/10 |
| | 17MEW7838 : 562290/11 |
| | 17MEW7838 : 562290/12 |
| | 17MEW7839 : 562290/14 |
| | 17MEW7839 : 562290/13 |
| | 17MEW7839 : 562290/15 |
| | 17MEW7839 : 562290/16 |
| | 17MEW7840 : 562290/17 |
| | 17MEW7840 : 562290/18 |
| | 17MEW7840 : 562290/19 |
| | 17MEW7840 : 562290/20 |
| | 17MEW7843 : 562290/26 |
| | 17MEW7843 : 562290/27 |
| | 17MEW7843 : 562290/28 |
| | 17MEW7843 : 562290/29 |
| | 17MEW7841 : 562290/21 |
| | 17MEW7841 : 562290/22 |
| | 17MEW7842 : 562290/25 |
| | 17MEW7841 : 562290/23 |
| | 17MEW7841 : 562290/24 |
| | 19AE3110 : 583659/01 |
| | 19AE4505 : 583667/01 |
| Tunisia | 6138 : 564401/01 |

| | |
|---|---|
| Environment Agency – Abu Dhabi, United Arab Emirates | 18AE445 : 565717/01<br>18AE10349 : 591851/02<br>20AE7901 : 591851/02 |
| San Diego Zoo Wildlife Alliance, USA | Loaned between registered scientific institutions (16US508 18A/9 to COSE GB039) |

All live sampling in Tunisia was undertaken by qualified veterinarians following a protocol approved after ethical review by the Marwell Wildlife Ethics Committee. Permission was granted by the Direction Générale des Forêts (Ministère de l'agriculture, des Ressources Hydrauliques et de la Pêche Maritime, Tunisie) for collection of samples in the field. Blood and tissue samples collected from zoological animals were excess to veterinary procedures.

**DNA extraction**

DNA was extracted from blood and tissue samples using either of the DNeasy Blood & Tissue kit (QIAGEN), QuickGene DNA whole blood kit S or tissue kit S (FUJIFILM Wako Chemicals Europe GmbH) following the manufacturers' protocols. Samples from AZA institutions were additional treated with RNase A as per Qiagen standard protocols for the tissue type. Faecal samples were extracted using the QIAmp DNA Stool Mini Kit (QIAGEN) or the Isohelix Xtreme DNA Kit (Cell Projects Ltd.) following the protocol described in (Werhahn et al., 2017).

**Mitochondrial DNA sequencing**

Species identification of all faecal samples was confirmed as addax by amplification of cytochrome b using the primers mcb398 (5'–TACCATGAGGACAAATATCATTCTG) and mcb869 (5'–CCTCCTAGTTTGTTAGGGATTGATCG) following Verma and Singh (2002). PCRs were carried out in 10 µL reactions containing 1x DreamTaq™ Hot Start PCR mastermix (ThermoFisher, contains 2.8 mM Mg++ and 280 µM of each dNTP), 2 µM of each primer and 1 µL of DNA. PCR thermocycling conditions were 95 °C for 5 mins; then 40 cycles of 95 °C for 30 sec, 50 °C for 30 sec, 72 °C for 60 sec; ending with 72 °C for 10 mins.

A 723 bp fragment of the mitochondrial (mtDNA) control region was amplified using the primers AdnewF (5'–GCTATAGCCCCACTATCAAC) and AdnewR (5'–GCGGGTTGCTGGTTTCACGC). The control region (also known as the D-Loop) was selected as this commonly shows variation within and between populations of a species (Avise et al. 1987PCR reactions were performed in 10 µL reactions containing 1.4X DreamTaq™ Hot Start PCR mastermix (contains 2.8 mM Mg++ and 280 µM of each dNTP), 2 µM of each primer and 1 µL of DNA. PCR thermocycling conditions were an initial denaturation of 4 mins at 94 °C; 10 cycles of 94 °C for 30s, 57 °C for 15s and 72 °C for 1min; 33 cycles of 95 °C for 30s, 54 °C 15s and 72 °C for 60s; with a final extension at 72 °C for 10 mins. PCR products were visualised by gel electrophoresis (1% agarose) and, when a single, clear band was detected.

The Adnew primers occasionally generated multiple PCR fragments, particularly when amplified from blood samples, which subsequently generated unreadable Sanger sequences. Nuclear copies of mitochondrial sequences, NUclear MiTochorndial sequences (NUMTs) are prevalent across eukaryotic genomes Hazkani-Covo, Zeller, and Martin (2010), including mammalian genomes Thalmann et al. (2004). In some species, the majority of the mitochondrial genome has been detected intact in NUMTS (e.g. humans - see Marshall and Parson (2021) and references within; felines Lopez et al. (1994)) The genome of the cow (Bos taurus) includes somewhere between 166 and 432 NUMT regions Calabrese et al. (2017). Separating organellar genomes from NUMTs in massively parallel sequencing data can present challenges (Maude et al. 2019), and whole genome resequencing data for scimitar-horned oryx (unpublished) and a divergent mitogenome published on Genbank (accession JN869311) indicates the high likelihood of large NUMTs in the scimitar-horned oryx and possibly Hippotraginae. It is therefore likely that the Adnew primers were co-amplifying organellar and NUMT control region sequences, and the lower mitochondrial to nuclear ratio in blood samples compared to tissue or faecal samples may have increased their prevalence.

Primers were designed to amplify the full-length control region sequence (tRNA-ProF 5'-CCCACTATCAACACCCAAAGC and tRNA-Phe 5'-AGCATTTTCAGTGCCTTGC). Single band amplifications were obtained for a handful of samples, but gel electrophoresis and sequencing indicated the continued presence of multiple bands for the majority of previously unsuccessful samples using the Adnew primers. This suggests that the full-length control region is present in a minimum of one nuclear copy.

Consequently, samples were amplified using the Adnew primers using a high-fidelity DNA polymerase, and the PCR fragment corresponding to the true mitochondrial sequence was gel-extracted. PCRs were carried out in 10µL reactions containing 1X Q5 high-fidelity DNA polymerase mastermix (NEB; contains 2 mM Mg++ and 200 µM of each dNTP), 0.5 µM of each primer and 1 µL of DNA (diluted 1:10). PCR thermocycling conditions were an initial denaturation of 4 mins at 98 °C; 10 cycles of 98 °C for 30s, 57 °C for 15s and 72 °C for 1min; 33 cycles of 95 °C for 30s, 54 °C 15s and 72 °C for 60s; with a final extension at 72 °C for 10 mins. All 10 µL of PCR product was run on a 1% agarose gel and extracted using the MinElute Gel Extraction kit (QIAGEN).

All mitochondrial DNA sequences were cleaned using 20U of Exonulease I (Thermo Scientific™) and 1U of FastAP (Thermo Scientific™) and were sequenced in both directions using BigDye Terminator v3.1 Cycle Sequencing Kit (Applied Biosystems) on a ABI 3730XL genetic analyser. Sequences were analysed using Geneious 2019 Prime (https://www.geneious.com).

**Mitochondrial DNA analysis**

Sequences were trimmed and quality checked by eye and using the heterozygote plugin (secondary peak height greater than 30%) to look for low quality bases. Sequences were aligned using the Geneious aligner and included an additional 12 published control region sequences (GenBank accessions: JN632591 (Hassanin et al. 2012), MZ474955-MZ475965 (Hempel et al. 2021)). Haplotypes were initially identified from an alignment by eye, then matched against a custom BLAST database in Geneious 2019 Prime. The GenBank sequence MZ474957 (museum sample from Sudan) was excluded due to the presence of two degenerate sites within the control region making haplotype determination unreliable; however both our control region data and the mitogenome analysis within Hempel et al. (2021) found that that MZ474957 clusters extremely closely with MZ474955 (museum sample from Sudan), suggesting it is not a novel haplotype. A string of 63 Ns was present in the MZ475964 (museum sample from Libya) sequence (see Suppl. Fig 1), encompassing an invariable region across all other haplotypes. Several methods for accounting for missing data and indels were tested (as below) and under all scenarios MZ475964 retained a unique control region haplotype, justifying its retention.

The presence of a 76 bp indel was detected within two haplotypes, with four polymorphic sites within the indel. and in both cases, the indel is an exact copy of the 76 nucleotides preceding it (see Suppl. Fig 1). It could be reliably and repeatedly amplified and unambiguously identified within multiple individuals (AC = 8 samples, L = 2 samples) from both wild collected faecal samples and captive collected blood samples, suggesting it is unlikely to be a NUMT. Small indels are also present within the haplotypes identified here (up to 4 bp in length), with only one being present across multiple haplotypes (a 4bp indel on haplotypes A, AC, F and AJ235310).

The large 76bp indel, in particular, presents a challenge for phylogenetic analyses and diversity estimates because its presence/absence represents a mutational event which, due to the high sequence similarity between the two indel sequences, are unlikely to represent two evolutionary events. However, analytical methods typically consider each nucleotide position independently, treating each polymorphism as an independent evolutionary event and sites containing a gap. Haplotype L is identical to haplotype P outwith the 76 bp indel, and therefore full exclusion of the indel omits evolutionarily informative site and complete deletion of indel sites is inappropriate for these data. We therefore assessed the impact of three methods of indel coding within the R package Haplotypes (Aktas 2020). Firstly, the simple indel coding (sic) method (Simmons & Ochoterena, 2000) treats an indel as a missing character but codes each separately following the simple indel coding method. Simulation studies suggested that simple index coding, and the related complex index coding, generate highly similar results which outperform other methods (Simmons, Müller, and Norton 2007). Secondly, the gaps are coded as a fifth state, under which each base is an independent evolutionary event. Finally, we assessed the impact of excluding the 76bp indel, imitating the standard treatment of small indels as missing data.

Statistical parsimony (TCS) haplotype networks (Templeton, Crandall, and Sing 1992) were selected as they can account for unobserved haplotypes (Paradis 2018). The addax data include haplotypes from wild and managed populations, as well as historical samples and we could therefore expect evolutionary intermediate haplotypes to be unsampled. As such, TCS networks were generated using the R pacakge Pegas (Paradis 2010) with the large 76bp indel re-coded as i) a single polymorphic site (imitating “sic” re-coding method), ii) multiple polymorphic sites (imitating the 5th state) and iii) complete deletion (imitating missing data). The PEGAS package does not perform recoding of indels as sic or fifth state, and performs complete exclusion of gaps. Although additional indels were present within the complete 723 bp (maximum of 5 bp length), only the 76 bp indel impacted topography of the haplotype networks, and so haplotype networks were generated for each of i) manual recoding of the whole 76 bp region as a single fifth state (imitating “sic”), ii) manual recoding each nucleotide as a fifth state, and iii) complete deletion of the 76 bp indel. All other indels were treated as default by PEGAS, which is complete deletion. Re-coding haplotypes as either “sic” or fifth state both recovered all 25 haplotypes. Under option iii) complete deletion of the indel, 24 haplotypes were recovered, and two haplotypes (L and P) were identical. Topologies for each method are shown below in Figure SM1.

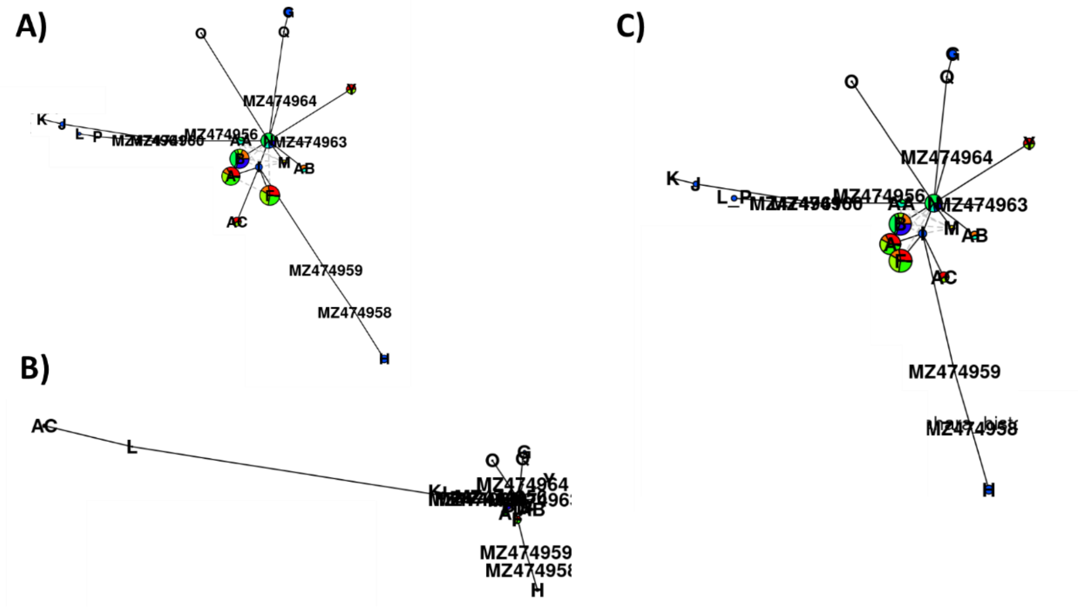


***Figure SM1.*** *Topologies of TCS networks for the 723 bp of addax control region sequences under the three scenarios for re-coding the 76bp indel. A) "sic" re-coding, B) 5$^{th}$ state re-coding and C) complete deletion of the 76 bp indel.*

Contemporary haplotypes were assigned alphabetical nomenclature but retained the GenBank accession number as nomenclature when detected in only within museum samples. The R package PEGAS (Paradis, 2010) was used to estimate nucleotide and haplotype diversity, as well as deviation from neutrality by estimating Tajima's D and calculating significance under a beta distribution (Paradis, 2020) with Bonferroni correction.

#### ddRAD library preparation and sequencing

DNA quality was assessed by gel electrophoresis using a 2% agarose gel and quantified using the Qubit dsDNA BR assay (Thermo Fisher Scientific), before normalisation to 7 ng/µL. A double digest restriction-site associated DNA (ddRAD) library was prepared following a modified Peterson et al (2012) protocol, as described in Bourgeois et al (2018). Briefly, the DNA was fragmented using the restriction enzymes SphI and SbfI and a unique pair of 5 or 7 bp barcodes were ligated to either end of the resulting DNA fragments of each sample. The pooled library was size selected between 400 and 700 bp using gel electrophoretic extraction, prior to PCR amplification to incorporate adapter sequences. Samples were sequenced across eight libraries, and positive controls were used to assess repeatability within and between libraries. Libraries were each sequenced (150 bp, paired-end) on a full lane of a HiSeq 2500/4000/X (Illumina, Inc).

**ddRAD data quality control and filtering**

The PROCESS_RADTAGS module of STACKS v2.52 (Rochette et al., 2019) was used to demultiplex the data, using default parameters to discard low-quality reads, filter for adapter sequences, remove reads with uncalled bases and trim the reads to 135 bp (as determined using FASTQC (Andrews, 2010)).

The scimitar-horned oryx (*Oryx dammah*) genome (REF-Humble2020) was used for reference-guided SNP identification. SNP calling was carried out using a custom SNAKEMAKE pipeline (Köster & Rahmann, 2012), available at [LINK github]. Briefly, ddRAD reads were mapped to the reference genome using BWA v0.7.17 (H. Li & Durbin, 2009), unmapped reads were filtered using SAMTOOLS v1.10 (Heng Li et al., 2009), and samples with fewer than 150,000 reads were excluded. STACKS v2.52 (Rochette et al., 2019) was used to call SNPs using the marukilow model with default parameters.

VCFTOOLS (Danecek et al., 2011) was used to filter low quality SNPs by first applying a minimum sequencing depth per individual genotype of 5 reads, a mean sequencing depth per SNP of 15 reads and a minor allele count of 3. A step-wise filtering scheme (as recommended by O'Leary et al., 2018) was then applied, first excluding SNPs with a genotyping rate <25%, then excluding individuals with a genotyping rate <50%, and finally retaining only SNPs with a genotyping rate of at least 50% in the remaining individuals.

Repeatability within and across libraries was assessed using positive control, before retaining only a single genotyping attempt for each repeated individual. Furthermore, relatedness was assessed within and between populations using KING (Manichaikul et al., 2010) to identify unintentionally duplicated (e.g. submitted under different IDs). All suspected duplicates (KING > 0.354) were corroborated by metadata and only the sample with the highest genotyping rate was retained. Linkage disequilibrium (LD) was estimated as $R^2$ in PLINK v1.9 (Chang et al., 2015), and we retained one from each pair of SNPs with an $R^2$ greater than 0.5.SNPs were then assessed for deviation from Hardy-Weinberg equilibrium (HWE) within each of the five main populations (EEP, SSP, EAD, AAZTunisia) using PLINK and were excluded if deviating in at least two populations. Lastly, all SNPs on the five main populations (EEP, SSP, EAD, AAZ and Tunisia) using PLINK and were excluded if deviating in at least two populations. Lastly, all SNPs on the sex chromosome (*O. dammah* chromosome 23) were removed and the SNP dataset was filtered using PLINK to include only those SNPs genotyped in 95% of individuals (1704 SNPs) – known as the relaxed LD SNP set.

A stringent LD SNP set was generated for analyses requiring minimum LD (admixture). The marker set was pruned to apply stringent LD filtering using PLINK v1.90 (Chang et al., 2015) to remove any SNP with an r-squared greater than 0.2 with any other SNP within a 50-SNP window using a 5-SNP sliding window, resulting in a dataset of 1073 SNPs.

**Literature cited**

Andrews, S. (2010). *FastQC: A Quality Control Tool for High Throughput Sequence Data*. http://www.bioinformatics.babraham.ac.uk/projects/fastqc/

Bourgeois, S., Senn, H., Kaden, J., Taggart, J. B., Ogden, R., Jeffery, K. J., Bunnefeld, N., Abernethy, K., & McEwing, R. (2018). Single-nucleotide polymorphism discovery and panel characterization in the African forest elephant. *Ecology and Evolution*, *4*, 2207–2217. https://doi.org/10.1002/ece3.3854

Chang, C. C., Chow, C. C., Tellier, L. C., Vattikuti, S., Purcell, S. M., & Lee, J. J. (2015). Second-generation PLINK: rising to the challenge of larger and richer datasets. *GigaScience*, *4*(1), 7. https://doi.org/10.1186/s13742-015-0047-8

Danecek, P., Auton, A., Abecasis, G., Albers, C. A., Banks, E., DePristo, M. A., Handsaker, R. E., Lunter, G., Marth, G. T., Sherry, S. T., McVean, G., & Durbin, R. (2011). The variant call format and VCFtools. *Bioinformatics*, *27*(15), 2156–2158. https://doi.org/10.1093/bioinformatics/btr330

Karesh, W. B., Smith, F., & Frazier-Taylor, H. (1987). A Remote Method for Obtaining Skin Biopsy Samples. *Conservation Biology*, *1*(3), 261–262. https://doi.org/10.1111/j.1523-1739.1987.tb00041.x

Köster, J., & Rahmann, S. (2012). Snakemake-a scalable bioinformatics workflow engine. *Bioinformatics*, *28*(19), 2520–2522. https://doi.org/10.1093/bioinformatics/bts480

Li, H., & Durbin, R. (2009). Fast and accurate short read alignment with Burrows-Wheeler transform. *Bioinformatics*, *25*(14), 1754–1760. https://doi.org/10.1093/bioinformatics/btp324

Li, Heng, Handsaker, B., Wysoker, A., Fennell, T., Ruan, J., Homer, N., Marth, G., Abecasis, G., & Durbin, R. (2009). The Sequence Alignment/Map format and SAMtools. *Bioinformatics*, *25*(16), 2078–2079. https://doi.org/10.1093/bioinformatics/btp352

Manichaikul, A., Mychaleckyj, J. C., Rich, S. S., Daly, K., Sale, M., & Chen, W. M. (2010). Robust relationship inference in genome-wide association studies. *Bioinformatics*, *26*(22), 2867–2873. https://doi.org/10.1093/bioinformatics/btq559

O'Leary, S. J., Puritz, J. B., Willis, S. C., Hollenbeck, C. M., & Portnoy, D. S. (2018). These aren't the loci you'e looking for: Principles of effective SNP filtering for molecular ecologists. *Molecular Ecology*, *27*(16), 3193–3206. https://doi.org/10.1111/mec.14792

Paradis, E. (2010). Pegas: An R package for population genetics with an integrated-modular approach. In *Bioinformatics* (Vol. 26, Issue 3, pp. 419–420). https://doi.org/10.1093/bioinformatics/btp696

Paradis, E. (2020). Population Genomics with R. In *Population Genomics with R*. Chapman and Hall/CRC. https://doi.org/10.1201/9780429466700

Peterson, B. K., Weber, J. N., Kay, E. H., Fisher, H. S., & Hoekstra, H. E. (2012). Double

Digest RADseq: An Inexpensive Method for De Novo SNP Discovery and Genotyping in Model and Non-Model Species. *PLoS ONE*, *7*(5), e37135. https://doi.org/10.1371/journal.pone.0037135

Rochette, N. C., Rivera-Colón, A. G., & Catchen, J. M. (2019). Stacks 2: Analytical methods for paired-end sequencing improve RADseq-based population genomics. *Molecular Ecology*, *28*(21), 4737–4754. https://doi.org/10.1111/mec.15253

Simmons, M. P., & Ochoterena, H. (2000). Gaps as Characters in Sequence-Based Phylogenetic Analyses on JSTOR. *Systematic Biology*, *49*(2), 369–381. https://www.jstor.org/stable/2585224

Verma, S. K., & Singh, L. (2002). Novel universal primers establish identity of an enormous number of animal species for forensic application. *Molecular Ecology Notes*, *3*(1), 220–222. https://doi.org/10.1046/j.1471-8286

Werhahn, G., Senn, H., Kaden, J., Joshi, J., Bhattarai, S., Kusi, N., Sillero-Zubiri, C., & Macdonald, D. W. (2017). Phylogenetic evidence for the ancient himalayan wolf: Towards a clarification of its taxonomic status based on genetic sampling from Western Nepal. *Royal Society Open Science*, *4*(6). https://doi.org/10.1098/rsos.170186
